# Tripartite interactions of PKA catalytic subunit and C-terminal domains of cardiac Ca^2+^ channel modulate its β-adrenergic regulation

**DOI:** 10.1101/2023.11.28.564875

**Authors:** Shimrit Oz, Tom Sharon, Suraj Subramaniam, Tamara Pallien, Moshe Katz, Vladimir Tsemakhovich, Debi Ranjan Tripathy, Giorgia Sasson, Orna Chomsky-Hecht, Leonid Vysochek, Maike Schulz, Claudia Fecher-Trost, Kerstin Zühlke, Daniela Bertinetti, Friedrich W. Herberg, Tal Keren-Raifman, Veit Flockerzi, Joel A. Hirsch, Enno Klussmann, Sharon Weiss, Nathan Dascal

## Abstract

The adrenergic nervous system augments cardiac contraction by increasing the activity of L-type voltage-gated Ca_V_1.2 channels. Dysregulation of this process is linked to severe cardiac dysfunctions. The signaling cascade involves activation of β-adrenergic receptors, elevation of cAMP levels, separation of protein kinase A (PKA) regulatory subunit (PKAR) from catalytic subunit (PKAC), and phosphorylation of the inhibitory protein Rad leading to increased Ca^2+^ influx. In cardiomyocytes, the core subunit of Ca_V_1.2 (α_1C_) exists in two forms: full-length (FL) or proteolytically processed (truncated), lacking the distal C-terminus (dCT). Specificity and efficiency in the cascade are believed to emanate from unique protein-protein interactions, such as anchoring PKA (via PKAR) to α_1C_ by A-kinase anchoring proteins (AKAPs). However, most AKAPs do not interact with the truncated α_1C_, and their role in βAR regulation of cardiac Ca_V_1.2 remains unclear. Here we show that PKAC, independently of PKAR or AKAPs, directly interacts with α_1C_ at two domains in α_1C_-CT: the proximal and distal C-terminal regulatory domains (PCRD and DCRD), which also interact with each other. Furthermore, we find that DCRD competes with PCRD and reduces its interaction with PKAC. The physiological consequences of these complex interactions are incompletely understood; our data suggest that they may fine-tune the βAR regulation of Ca_V_1.2. We propose that the newly discovered interactions take part in governing colocalization of regulatory proteins within the βAR-Ca_V_1.2 multimolecular signaling complexes in cardiomyocytes.

## Introduction

Ca^2+^ influx via L-type voltage-gated Ca^2+^ channels (Ca_V_1.2) underlies cardiac excitation-contraction coupling (1). Ca_V_1.2 channels are prominently modulated by epinephrine and the sympathetic nervous system, underlying much of the physiological regulation of heartbeat and the “fight-or-flight” response, via the activation of β-adrenergic receptors (βAR), β1AR and β2AR (2). The ensuing signaling cascade leads to elevated cAMP levels, activation of protein kinase A (PKA) by the disassembly of the PKA holoenzyme into catalytic (PKAC) and regulatory (PKAR) subunits (3,4) (Fig. 1), and culminates in the enhancement of Ca_V_1.2 channel currents. In cardiomyocytes (CM) much of this signaling occurs within precise micro/nanodomains (signalosomes) that have been proposed to include the βARs, G proteins, adenylyl cyclase, PKA, Ca_V_1.2 and additional interacting proteins (5-9). Dysregulation of this signaling cascade may result in heart failure, certain arrhythmias, such as Catecholaminergic Polymorphic Ventricular Tachycardia (CPVT) (10). β1ARs are located mostly at the crest and signal globally; β2ARs interact and co-localize with Ca_V_1.2 at the t-tubules, and signal locally (5,11-15). Despite decades of intense research (16), crucial aspects of this regulation, such as the spatiotemporal organization of the signaling complexes and the target(s) of PKA phosphorylation, remain unclear or debatable (17-19). Here we report a novel direct interaction of PKAC with two regulatory segments of Ca_V_1.2, PCRD and DCRD. We propose that this complex tripartite interaction plays a previously unrecognized role in organizing the signaling complex and fine-tuning the βAR effect.

**Figure 1.**
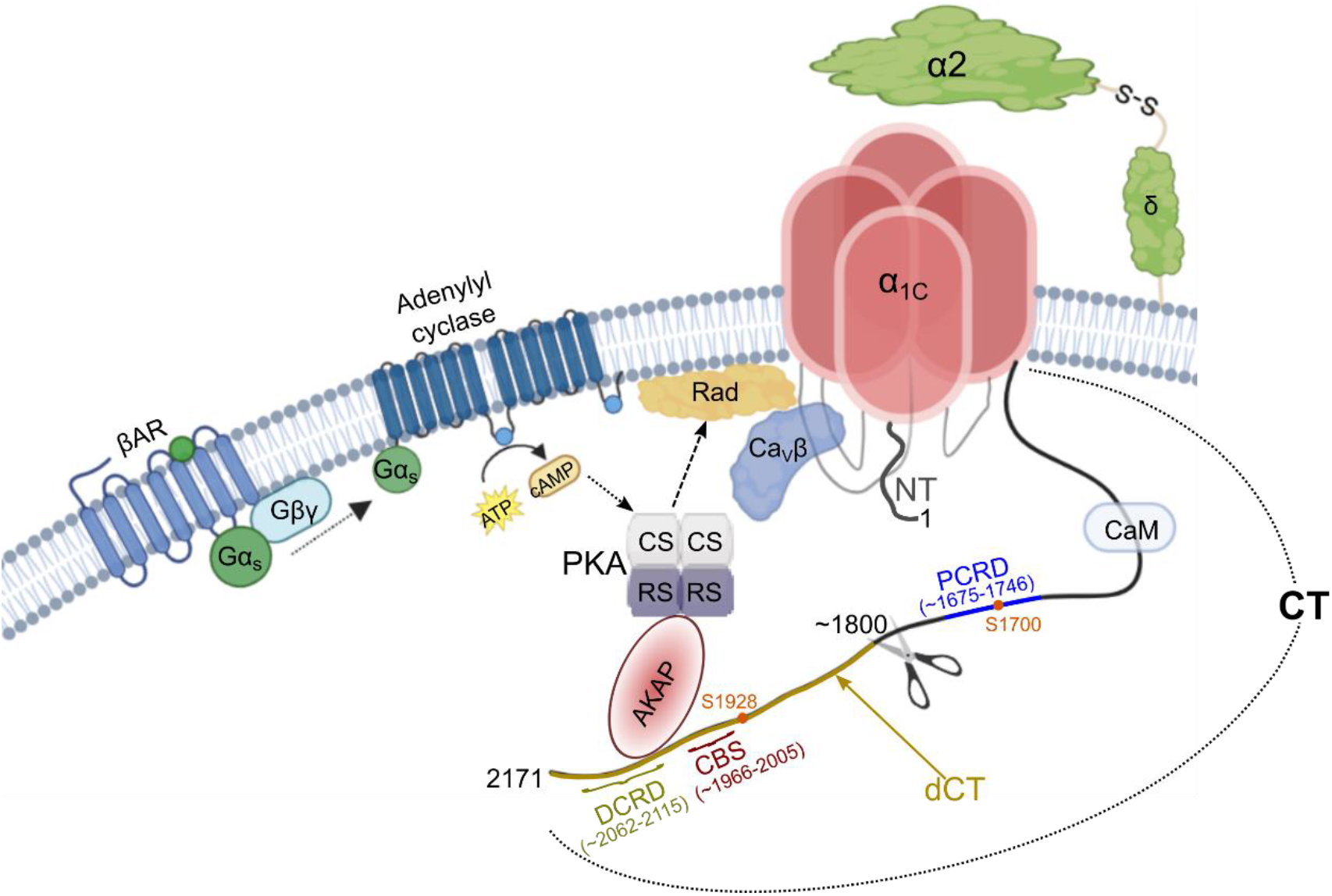
The current concept of the signaling pathway underlying the β-adrenergic regulation of Ca_V_1.2 in cardiomyocytes. The cascade is initiated by the binding of an agonist (e.g. epinephrine or norepinephrine; depicted as a small green circle) to a β-adrenergic receptor (βAR) which, in turn, activates the G protein G_s_ by catalyzing GDP-GTP exchange at the Gα_s_ subunit followed by separation of the latter from the Gβγ subunit. The GTP-bound Gα_s_, alone or in concert with Gβγ, activates adenylyl cyclase (AC), promoting the conversion of ATP to cAMP. cAMP activates PKA, causing the dissociation of its regulatory (RS) from catalytic (CS) subunits. CS phosphorylates several targets including α_1C_ and Ca_V_β subunits, but the most important target in Ca_V_1.2 regulation appears to be the Rad protein. Phosphorylation of Rad removes the constitutive inhibition that it exerts upon channel activity, through separation of Rad from Ca_V_β. Before activation, the PKA holoenzyme is thought to be associated with the α_1C_ subunit via an AKAP protein that strongly binds PKAR (the binding main sites for most AKAPs are in the dCT of α_1C_). The scheme also shows the auxiliary subunits of Ca_V_1.2 and the cytosolic domain of α_1C_: N-terminus (NT), C-terminus (CT) (with dCT colored dark mustard), and intracellular loops (shown in light gray). Also indicated are the approximate CT proteolytic cleavage site (scissors), major PKA phosphorylation sites in the CT (S1700 and S1928), and the DCRD, CBS, and PCRD domains.

Ca_V_1.2 channels comprise the pore-forming subunit α_1C_, intracellular Ca_V_β and calmodulin (CaM), and extracellular α2δ (20-22). Much of the cardiac α_1C_ is post-translationally cleaved at the C-terminus (CT), around amino acid (a.a.) 1800-1820, to produce the truncated α_1C_ protein and the cleaved distal CT (α_1C_-dCT) (23-26) (Fig. 1); however, full-length (FL) α_1C_ protein is also present (24,25,27).

The α_1C_-dCT is an important regulator of Ca_V_1.2. In the full-length channel it acts as an inhibitory module that reduces the channel’s open probability and voltage sensitivity (28,29). It also serves as a binding hub for signaling proteins, such as the protein phosphatase PP2A, several A-kinase anchoring proteins (AKAPs), β2AR, and others (15,27,30-33), and was suggested to take part in PKA regulation of the channel (2,24,34). An unknown fraction of the cleaved α_1C_-dCT translocates to the nucleus and acts as a transcription regulator for α_1C_ and other proteins (35,36), and was also proposed to remain bound to, and inhibit, the truncated channel (24). In neurons, dCT cleavage is Ca^2+^-dependent and catalyzed by calpain (37). In contrast, in cardiomyocytes, neither the specific protease nor the proportion of the channel-bound vs. nuclear α_1C_-dCT, or the fraction of FL and truncated α_1C_ are known; the latter may be species, organ, and age-dependent (23,38,39).

Two separate regulatory domains within α_1C_-CT were previously identified (24): the sequence proximal to the truncation (PCRD: Proximal C-terminus Regulatory Domain; a.a. 1675- 1746) and a sequence distal to the truncation (DCRD: Distal C-terminus Regulatory Domain) (Fig. 1). The DCRD, either as a part of the FL channel or after cleavage, inhibits the channel due to its interaction with PCRD (24). Interestingly, proximally to DCRD, a novel Ca_V_β-binding site in the α_1C_- dCT, termed C-terminal β-Binding Site (CBS), has been recently identified (40) (Fig. 1).

Spatial and functional organization of βAR/PKA/Ca_V_1.2 signaling involves AKAPs, scaffolding proteins orchestrating the assembly of cell compartment-specific macromolecular complexes (27,41-46). In the classical scheme, the PKA holoenzyme is docked to α_1C_-dCT indirectly, via an AKAP that possesses separate binding sites for PKAR and α_1C_-dCT and the plasma membrane (Fig. 1). Binding of cAMP to PKAR releases PKAR from PKAC, which then phosphorylates the target protein (33,47-49). However, it is unclear how important AKAPs are for the βAR regulation of cardiac Ca_V_1.2. Initial reports suggested a central role of AKAP5 (AKAP79/150) or AKAP7 (AKAP15/18) in βAR regulation of Ca_V_1.2 in cardiomyocytes (48,49), yet, genetic deletion of AKAP5 and AKAP7 did not significantly attenuate the βAR up-regulation of Ca_V_1.2 in cardiomyocytes (50,51), though deletion of another AKAP, Cypher/ZASP, weakened the regulation (32,52). Moreover, the presence of dCT that contains the main binding site for most AKAPs is not essential for PKA regulation, since heterologously expressed Ca_V_1.2, either full-length or dCT-truncated, is robustly regulated by activation of PKA and βARs without AKAP coexpression (53,54). Thus, while AKAP anchoring of PKA may be crucial for Ca_V_1.2 regulation in some organs (19), it does not appear essential in cardiac muscle.

Importantly, it is unclear whether full length and truncated α_1C_ forms of Ca_V_1.2 are differently regulated by βARs and PKA. The most prominent PKA phosphorylation sites in α_1C_, Ser1700 and Ser 1928, are located in PCRD and in the cleaved α_1C_-dCT, respectively. Direct phosphorylation of one of these sites has long been believed to underlie the PKA regulation of Ca_V_1.2 (2). This appears to hold true for the phosphorylation of Ser1928 in neurons and smooth muscle (55-58), and it may be involved in channel oligomerization upon βAR activation (59). However, it is not the case in the heart, as shown by studies in cellular systems and genetically engineered mice (17,18,25,60,61). Recent seminal studies (53,62-64) identified phosphorylation of Rad as an essential step in βAR-Ca_V_1.2 regulation in the heart (Fig. 1). Rad constitutively inhibits cardiac Ca_V_1.2, mainly via an interaction with the β subunit (65-69) (Fig. 1). PKA phosphorylation of Rad removes it from the plasma membrane, disrupts Ca_V_1.2-Rad proximity and Ca_V_β-Rad interaction, and alleviates Rad’s inhibitory effect on Ca_V_1.2 (53,62,70,71). Recent studies decisively demonstrated that coexpression of Rad is essential for the reconstitution of the adrenergic regulation of Ca_V_1.2 in heterologous expression systems (53,54). Nevertheless, a regulatory role for α_1C_ phosphorylation has not been ruled out, since mice with mutated phosphorylation sites on α_1C_ exhibit impaired baseline regulation of cardiac contractility (72,73). It also remains unclear whether the autoinhibition of Ca_V_1.2 function by the α_1C_-dCT domain, and its interaction with the proximal CT, play a role in PKA regulation, as suggested (2,74,75).

Given the robust βAR regulation of the truncated Ca_V_1.2 (54), lacking the key AKAP-binding sites, we hypothesized that there may be an AKAP-independent mechanism to ensure PKA proximity to α_1C_ following adrenergic stimulation. We found that PKAC co-precipitates with dCT-truncated α_1C_ (α_1C_Δ1821), independently of PKAR. We then mapped the interaction sites to both PCRD and DCRD and demonstrated their direct interaction with PKAC and with each other. DCRD competes with PCRD and reduces its interaction with PKAC. Our results suggest unique interaction sites for PKA on α_1C_-PCRD, present in both FL and truncated channels, and in the dCT (only in FL channels). These complex, AKAP-independent, interactions likely regulate the efficacy of the signaling cascade, resulting in quantitative differences in βAR modulation of FL and truncated channels.

## Results

### Protein kinase A catalytic subunit and α_1C_Δ1821 co-immunoprecipitate independently of PKA-regulatory subunit

To examine the possibility that PKA interacts directly with a truncated α_1C_ lacking the AKAP-binding α_1C_-dCT, we have generated a HEK293 cell line stably expressing tetracycline-inducible mouse α_1C_Δ1821 (Fig. S1A, B). We found that these cells also express endogenous Ca_V_β subunits (Fig. S1C), which most likely contributes to the formation of functional channels. Induced expression of α_1C_Δ1821 resulted in calcium channel currents that showed a typical current-voltage relationship (Fig. S1D). Next, we transiently transfected the α_1C_Δ1821 HEK293 cells with YFP-fused PKAC (PKAC-YFP) and performed immunoprecipitation (IP) of α_1C_ (Fig. 2A). IP with the α_1C_ antibody, but not Na_V_1.1 sodium channel antibody, specifically precipitated α_1C_Δ1821. Importantly, we observed co-immunoprecipitation (co-IP) of PKAC-YFP with α_1C_Δ1821. PKAC-YFP was detected by the PKAC antibody at ∼70 kDa only in cells that expressed both α_1C_ Δ1821 and PKAC-YFP and when the IP was done with the α_1C_ antibody, but not in the control IP with the Na_V_1.1 antibody (Fig. 2A, right panel).

**Figure 2.**
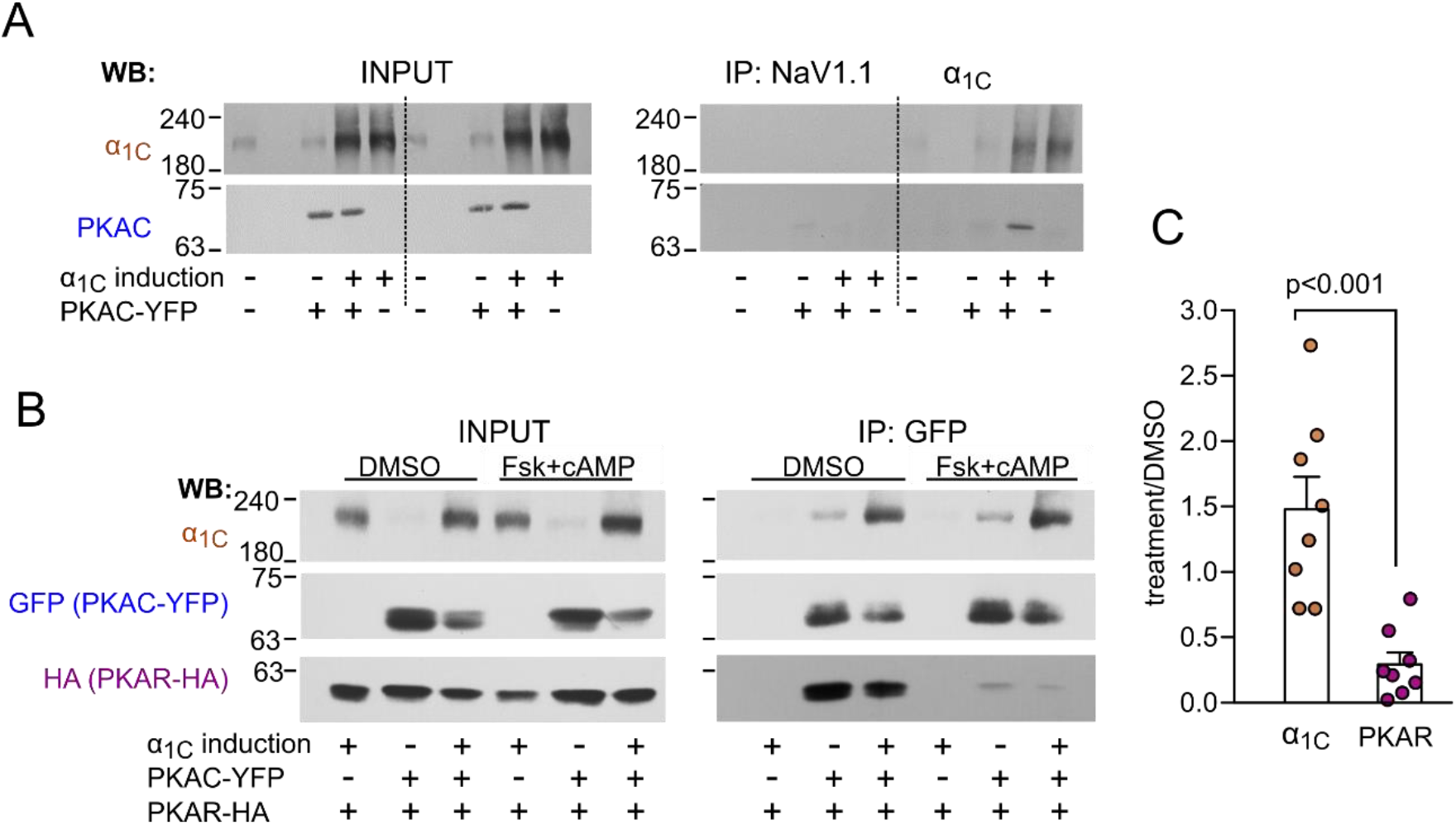
α_1C_Δ1821 and PKAC co-immunoprecipitate independently of PKAR. *A, left panel:* Tetracycline administration induced α_1C_ expression in HEK293 cells stably transfected with α_1C_Δ1821, with or without transiently expressed PKAC-YFP (marked as input). *Right panel*: α_1C_ antibody, but not sodium channel subunit antibody, Na_V_1.1, specifically enriched α_1C_Δ1821 and co-precipitated transiently expressed PKAC-YFP (detected by PKAC antibody) only in cells that co-express α_1C_Δ1821 and PKAC-YFP. *B, left panel:* Tetracycline administration induced α_1C_ expression, with or without transiently expressed PKAC-YFP and PKAR-HA (marked as input). *Right panel*: GFP antibody specifically enriched PKAC-YFP and co-precipitated α_1C_Δ1821 and PKAR-HA. Addition of forskolin and cAMP, but not DMSO, lowered the amount of co-precipitated PKAR, but did not affect co-precipitated α_1C_Δ1821. *C,* PKA activators enhanced the interaction of PKAC with α_1C_Δ1821 (1.47 ±0.25, n=8) and reduced its interaction with PKAR (0.29 ±0.09, n=8). Y axis shows normalized band intensity, treatment/control (without PKA activators, with DMSO only).

In a complementary experiment (Fig. 2B), transiently expressed PKAC-YFP (precipitated with the GFP antibody) co-precipitated α_1C_Δ1821 and the transiently expressed HA-tagged PKA-regulatory subunit (PKAR-HA, detected with anti HA antibody). PKA activators (forskolin and/or cAMP) considerably decreased the amount of PKAR co-precipitated with PKAC, as expected. In contrast, the amount of co-precipitated α_1C_Δ1821 did not decrease but actually tended to increase following this treatment (Fig. 2B, right panel). Thus, PKA activation by forskolin and/or cAMP increased the amount of α_1C_Δ1821 (1.47 ±0.25, n=8), and reduced the amount of PKAR interacting with PKAC (0.29 ±0.09, n=8) compared to control (Fig. 2C). Taken together, these results suggest that PKAC directly interacts with α_1C_Δ1821, independently of its interaction with PKAR or the presence of α_1C_-dCT.

### Mapping PKAC and α_1C_-CT interaction sites

We used scanning peptide array analysis to explore and identify interaction sites in α_1C_ and PKAC. Since the initial part of the NT and residues within the CT, but not the cytosolic loops 1-3 connecting the channel’s repeating domains, have been implicated in PKA regulation of Ca_V_1.2 (49,55,58,75,76), we prepared an array of 25-mer spot-immobilized overlapping peptides covering the first half of human and rabbit NT and the whole CT domain of rabbit α_1C_, excluding the specific calmodulin-binding IQ motif. We overlaid the membrane with purified His-PKAC protein and probed it with PKAC antibody (Fig. S2). We detected moderate binding to the initial 25 a.a. of the human NT (Fig. S2, spot E16); this binding was enhanced by alanine mutations of most a.a. residues in this peptide (Fig. S2, spot E16). However, no binding was detected in the homologous initial segment of the rabbit α_1C_ (88% identity), which was the model channel used in this and most previous studies, and which is robustly regulated by PKA in model reconstitution systems (53,54,62). We did not observe significant PKAC binding sites in any of the other peptides corresponding to the NT domain of rabbit α_1C_.

In contrast, we reproducibly observed strong interaction (black spots) of PKAC with three domains in α_1C_-CT: K1672-G1731, V1737-E1776 and L2072-L2126 (Fig. 3A, B; Fig. S2). Interestingly, the two main interaction domains partially overlap PCRD (Q1675 -S1746) and DCRD (S2062 - G2115), the two domains previously proposed to interact with each other, underlying the auto-inhibitory action of the dCT (24). Notably, PCRD contains the PKA recognition sequence ^1696^RRxxS^1700^ with the well characterized PKA phosphorylation site S^1700^ (2).

**Figure 3.**
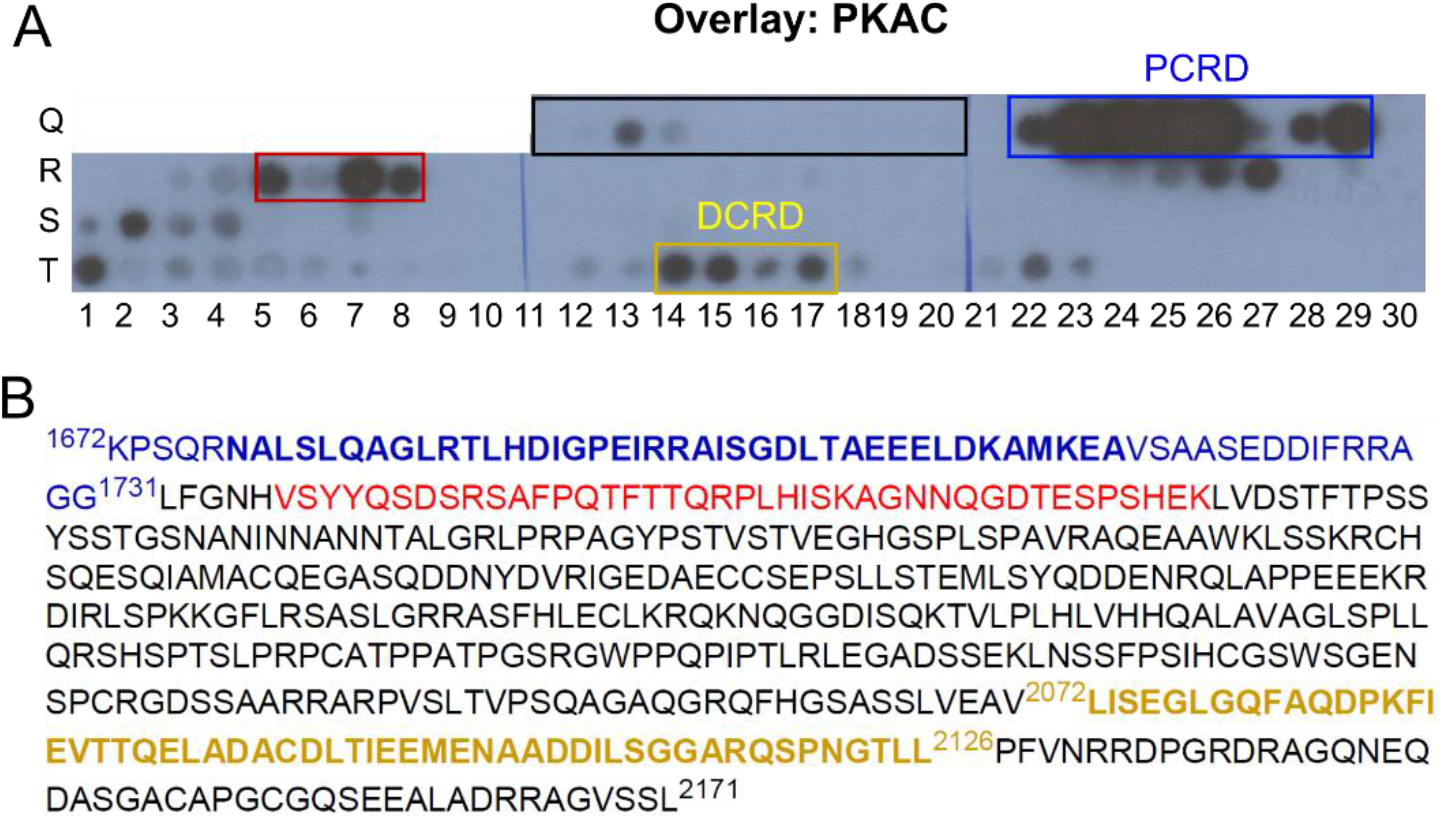
Peptide array reveals PKAC interaction sites within α1C Δ1821-CT. *A,* An array of 25-mer overlapping peptides with a 5 a.a. shift from α_1C_-CT segment, immobilized as spots on a membrane, were overlaid with purified His-PKAC and probed with PKAC antibody. Shown is the part of the membrane presenting the CT region between a.a. 1576-1647 (framed in black box) and a.a. 1672-2171 (the rest of the array). Within this segment, cysteines were replaced with serines to avoid the formation of disulfide bridges. See Fig. S2 for the image of the full array and additional details. The overlay reveals three main interaction domains on α1C-CT: K1672-G1731 (PCRD; framed in blue box), V1737-E1776 (red box) and L2072-L2126 (DCRD; dark yellow box). *B,* Amino acid sequence of the interaction domains; color code same as boxes in A.

To further corroborate the interaction between PKAC and PCRD or DCRD segments of α_1C_, we performed a complementary scan of a PKAC peptide arrays with purified PCRD (G1671-T1751) and DCRD (R2058-S2120). We prepared DNA constructs for bacterial expression of His-DRCD-Myc and His-SUMO-PCRD. SUMO protein was fused to PCRD to improve expression and stability of the protein). Both proteins were expressed in *E. coli* and purified. His-DCRD-Myc (DCRD in brief) showed a strong protein band of the predicted molecular mass (11.5 KDa; Fig. S3). The purified His-SUMO-PCRD, appeared in two distinct molecular sizes that were detected by the SUMO antibody, indicating that the protein is partially truncated at its C-terminus. The full-length ∼25 kDa protein (“peak 2”, Fig. S4 A-C) tended to degrade during the experiment. The major protein fraction (“peak 3”, Fig. S4 A-C) contained a single main protein product running at ∼17 kDa, as shown by SDS PAGE and HPLC analysis (Fig. S4D). We have termed this protein His-SUMO-PCRD_trunc_, or PCRD_trunc_ for brevity. Further analysis of PCRD_trunc_ by mass spectrometry (Fig. S4E) confirmed that it extends to Y^1739^ and thus includes the main putative PKAC-binding, PCRD-overlapping segment detected in the peptide array (Fig. 3). It also contains the aforementioned PKA recognition sequence that is expected to bind PKAC. Importantly, this segment in PCRD is homologous to known PKA-binding pseudosubstrate sites found in PKAR and the specific PKA inhibitor, PKI (Fig. S4F) (77,78). The peptide array indicates a smaller putative PKAC binding site downstream from PCRD_trunc_ (labeled by red frame in Fig. 3A and red letters in Fig. 3B); we have not further studieds this putative interaction site.

We scanned the array of 25-mer overlapping peptides covering the full-length of PKAC, using overlays with DCRD (detected by Myc antibody) or PCRD_trunc_ (detected by His antibody). The overlay experiments suggested two surfaces on PKAC that interact with PCRD; one of these also interacted with DCRD (Fig. 4C). The binding sites partially overlap with RyR2 binding sites on PKAC (79). The array results also indicated possible additional shorter DRCD-binding segments, but they were not consistent between replicates. Overall, these results strengthen the notion that PKAC directly interacts with PCRD and DCRD.

**Figure 4.**
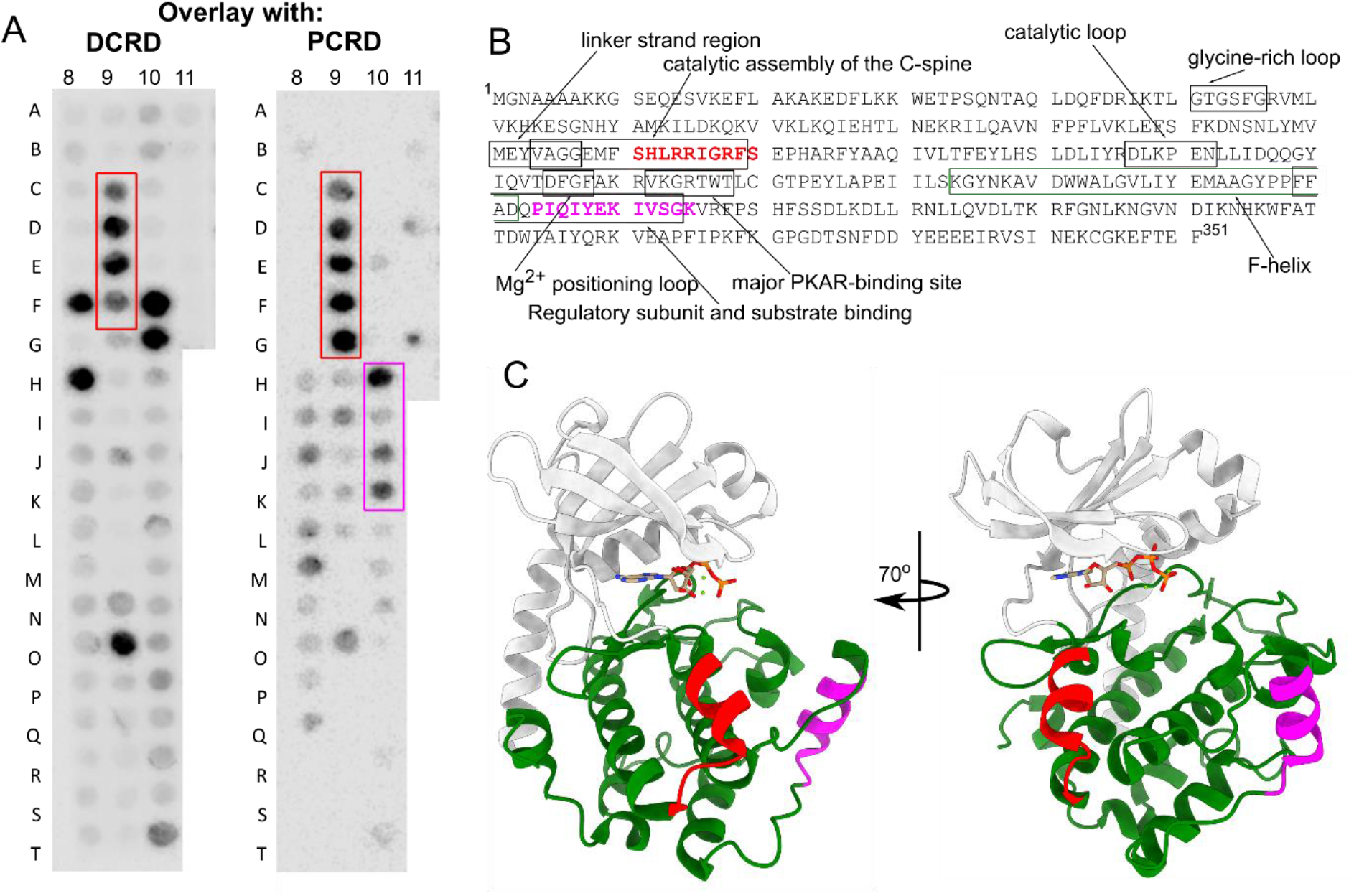
Peptide array reveals interaction sites of PCRD and DCRD with PKAC. *A*, An array of 25- mer overlapping peptides, with a 5 a.a. shift, of PKAC immobilized as spots on a membrane were overlaid with purified His-DCRD-Myc (left) and His-SUMO-PCRD (right). The overlay reveals a common DCRD/PCRD binding site (framed in red) and an additional PCRD unique site (magenta) on PKAC. Shown are representative results of three independent experiments. *B*, PCRD and DCRD binding sites on PKAC are located within solvent exposed helices. *C*, Side view (left) and a 70° rotated view (right) cartoons of human PKAC protein (PDB: 4WB5 (99)). The PKI peptide and amino acids 311-350 of this structure were removed for clarity. An ATP molecule with 2 metal ions is depicted as sticks in the catalytic site. The PKAC N-terminal domain is colored gray, the C-terminal domain green, the common PCRD/DCRD binding site red, and the unique PCRD binding site magenta.

### PCRD and DCRD directly interact with PKAC in a concentration-dependent manner

To further characterize the direct interaction between PKAC and the PCRD and DCRD domains of α_1C_, we performed pull-down experiments using purified GST-PKAC, PCRD_trunc_, and DCRD proteins. Western blots of the proteins eluted from Ni-NTA beads indicated that both PCRD and DCRD interact with PKAC in a concentration-dependent manner (Fig. 5). Control experiments verified that purified His-SUMO alone did not bind PKAC (Fig. S5). The binding data could be fitted to Hill’s equation with an apparent dissociation constant below 1 µM for both proteins. Since the pulldown procedure involves multiple steps and the local concentration of proteins attached to Ni-NTA beads is unknown, the apparent sub-micromolar affinity estimates may not reflect the actual affinity.

**Figure 5.**
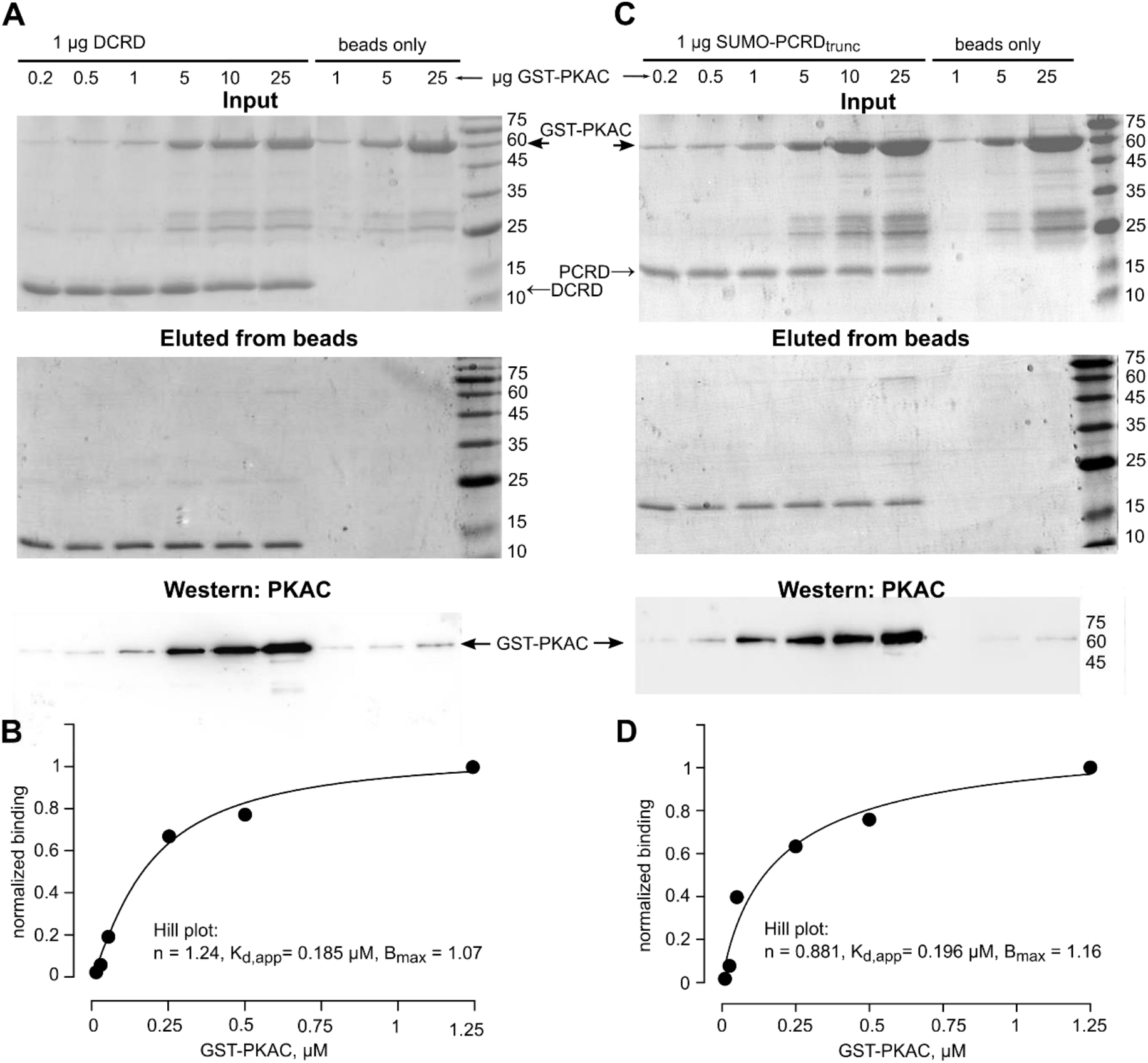
Dose dependent binding of GST-PKAC to DCRD and PCRD_trunc_. 1 µg of purified DCRD (A, B) or PCRD_trunc_ (C, D) were used to pull down the indicated amounts of GST-PKAC on Ni-NTA beads. The reaction volume was 300 µl. *A*, DCRD/PKAC interaction. Upper gel: input (Coomassie staining of a separate gel with exactly the same amounts of proteins as for pull-down reaction shown in the middle gel). Middle gel: elution from beads (Coomassie). Lower image: Western blot of the eluted proteins, with PKAC antibody. *B*, quantitation of the binding curve. Data were analyzed with ImageJ. Intensities of GST-PKAC binding were normalized to 25 µg GST-PKAC (lane 6). Data were fitted to a Hill equation in the form f=B_max_*[PKA]^n^/([PKA]^n^+K_d,app_^n^), where B_max_ is maximal binding, [PKA] is GST-PKAC concentration, K_d,app_ is the apparent dissociation coefficient, and n is Hill coefficient. The parameters of the fit are shown in the inset. Representative of 4 experiments. *C*, PCRD_trunc_/PKAC interaction (same presentation as in A). Upper gel: input (Coomassie); middle gel: elution from beads (Coomassie); lower image: Western blot of the eluted proteins, with PKA antibody. *D*, quantitation of the binding curve. Representative of 3 experiments.

To obtain a better estimate of the thermodynamic parameters of PKAC interaction with PCRD_trunc_ we used isothermal titration calorimetry (ITC) (Fig. 6). The ITC confirmed the concentration-dependent and saturable binding between PKAC and PCRD_trunc_ with a K_D_ of 1.4 µM. Interestingly, the estimated PCRD/PKAC molar ratio was 1.73, supporting the possibility of two PCRD binding sites in PKAC as suggested by the peptide array scan (Fig. 4).

**Figure 6.**
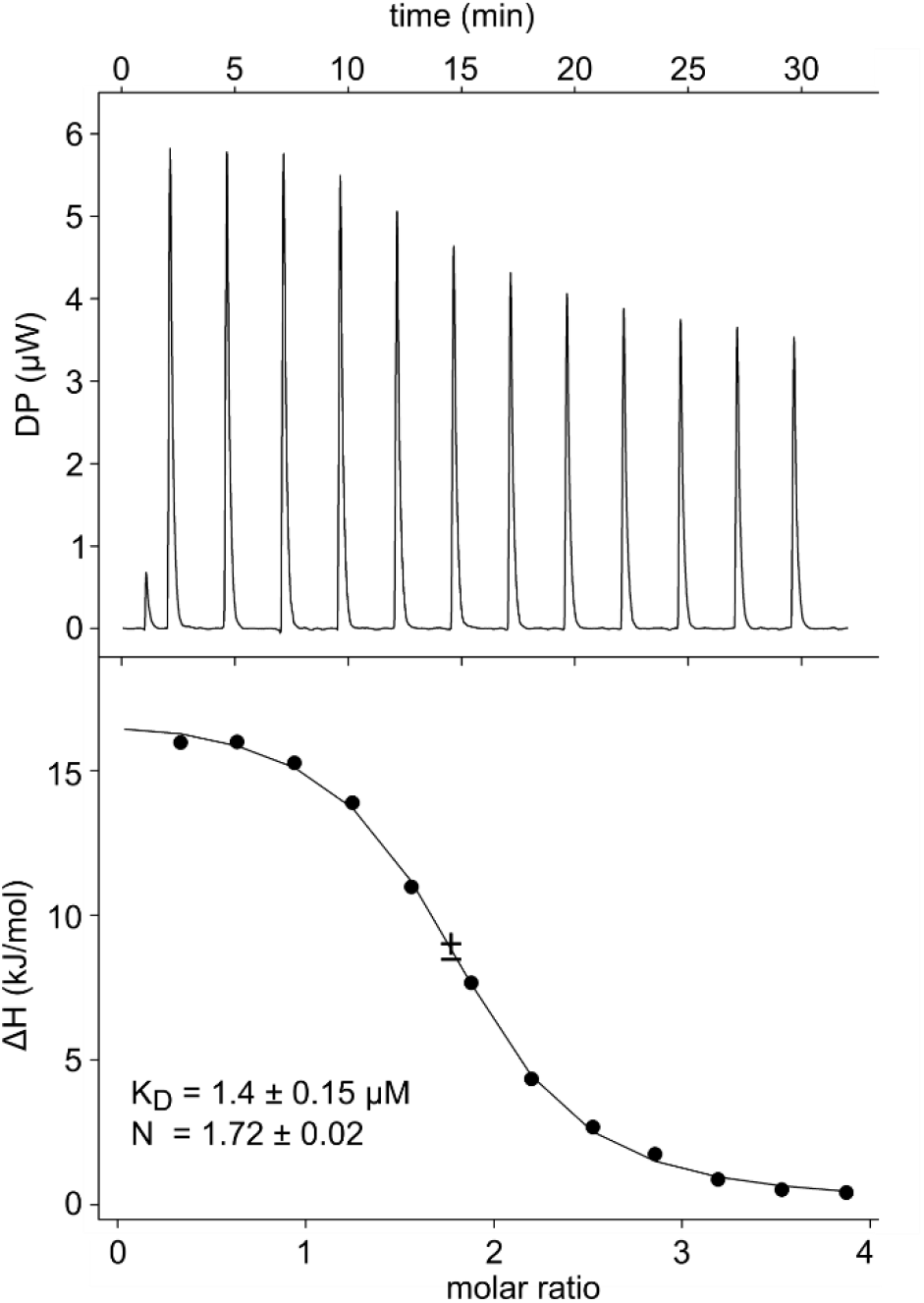
Calorimetric measurement for the titration of His-SUMO-PCRD into His-PKAC. In the top panel, data are shown as an injection profile of His-SUMO-PCRD [syringe; 400 µM] into His-PKAC (cell). The power output in DP (μW) was measured as a function of time in minutes. The heat of dilution of buffer and both proteins were subtracted and the area under the curve was integrated to generate the points that represent heat exchange in kJ/mole and plotted against the His-SUMO-PCRD to His-PKAC molar ratio for each injection, shown in the bottom panel. The solid line represents the best-fit curve for the data. Inset: thermodynamic parameters describing the fit, where N is the stoichiometry between PCRD and PKAC. The other thermodynamic parameters were: ΔG = -9.0 cal/mol; ΔH = 4.1 ± 0.77 cal/mol; -TΔS = -12.1 cal/mol. Representative of two experiments.

### Tripartite interaction between PKAC, PCRD and DCRD

PCRD and DCRD have been proposed to interact, presumably by forming salt bridges between R1696/R1697 (PCRD) and E2103/E2106/D2110 (DCRD) (24). Using synthetic fluorescein-labeled, 30- or 44-mer peptides, PCRD30 and PCRD44 (Fig. 7A), we found that purified recombinant His-PKAC and His-DCRD, but not PKAR (we used PKA-RII), bound the PCRD30 and PCRD44 peptides in a pull-down assay (Fig. 7B). The extent of binding to PKAC was similar for PCRD30 and PCRD44, suggesting that the distal part of PCRD44 (the last 14 a.a.) is not crucial for this interaction. This is the first experimental demonstration of a direct interaction between PCRD and DCRD of Ca_V_1.2.

**Figure 7.**
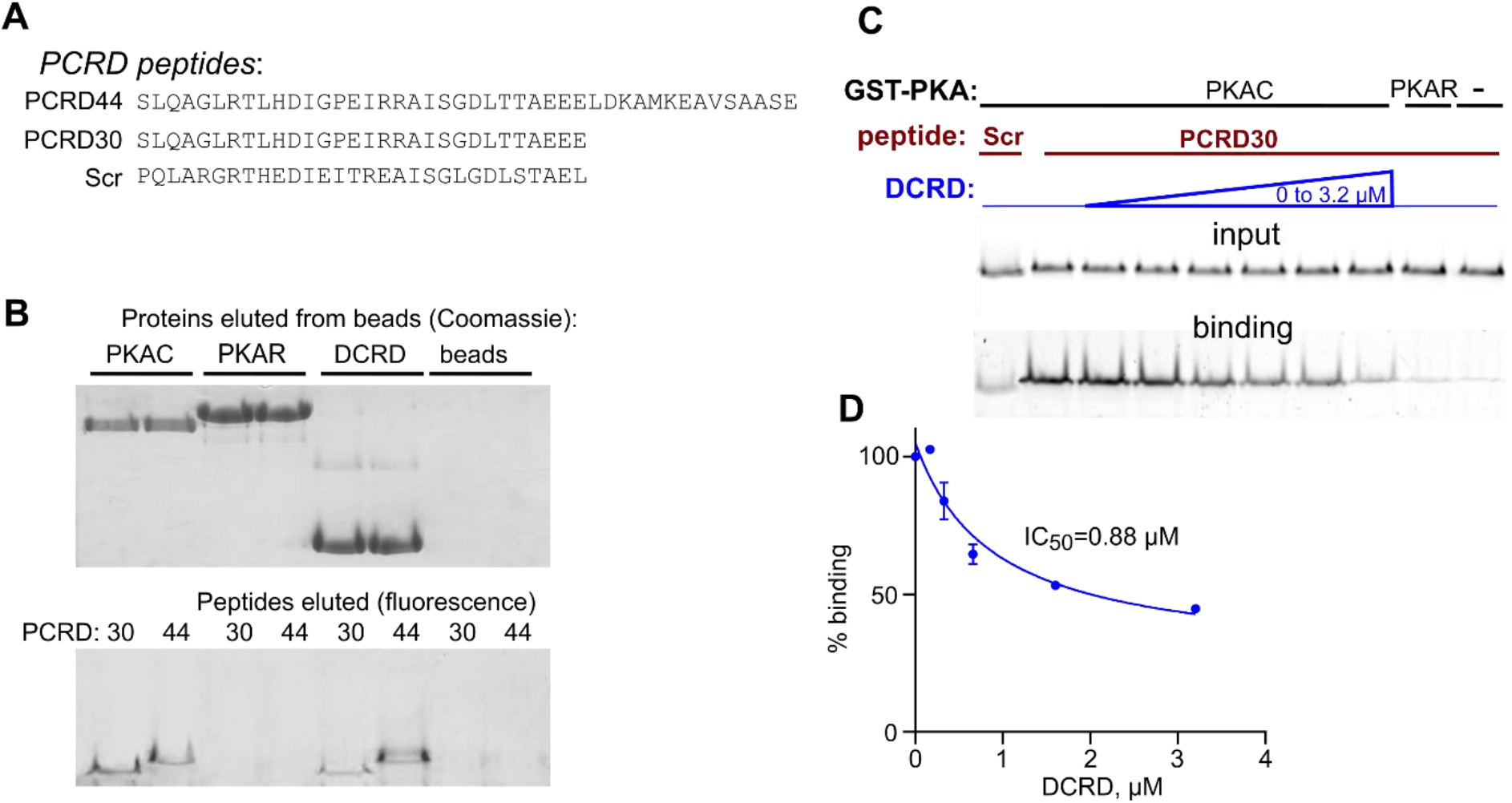
PKAC and DCRD, but not PKA-RIIβ, bind the PCRD peptides. *A*, Fluorescein-labeled peptides used were PCRD44 and PCRD30. Scrambled (Scr) peptide was used as a negative control. *B*, His-PKAC and His-DCRD, but not His-PKA-RIIb, bind the PCRD44 and PCRD30. The binding reaction was performed with 10 μg of each protein and peptide. Proteins were eluted from Ni-NTA beads and separated on a 16.5% polyacrylamide gel with Tricine buffer (1% SDS). SDS was not present in the sample buffer. Upper, Coomassie staining; lower, imaging of fluorescein-labeled PCRD peptides. *C*, DCRD impedes the PKAC/PCRD interaction. Binding reaction was performed with 250 nM GST-PKAC or GST-PKA-RIIb, 2.5 μM PCRD peptides, and increasing concentrations of His-DCRD (from 0 to 3.2 µM). Pull-down was done with glutathion affinity beads. *D*, summary of 3 experiments as in C. Each point shows mean±SEM. To construct the binding curve, data were fitted to a standard binding isotherm in the form [% of control=min + (max-min)/(1+(X/IC_50_))], yielding a half-inhibition concentration (IC_50_) of 0.88 µM, maximal (max) binding of 105.2%, minimal (min) binding level of 25.6%.

The confirmation of direct interaction between DCRD and PCRD, both of which bind PKAC at the same ∼30-a.a. segment (see Fig. 4), raised the possibility of competition between PKAC and DCRD for binding to PCRD. We examined the binding between GST-PKAC and PCRD30 in the presence of increasing concentrations of His-DCRD. GST-PKA-RII served as a negative control. Addition of DCRD resulted in decreased binding between PKAC and PCRD30, with half-inhibition concentration (IC_50_) of 0.88 µM (Fig. 7 C, D). DCRD did not cause a complete dissociation of PCRD30 from PKAC, leaving a significant PKAC-bound fraction of PCRD30 peptide (∼25% at saturating DCRD concentrations, as estimated from the results of fitting of the binding curve; Fig. 7D). These results suggest a tripartite interaction within the PKAC – PCRD – DCRD complex that includes partially overlapping common binding sites.

To address the potential functional consequences of the triple PKA-PCRD-DRCD interaction, we first tested the hypothesis that the interaction with PCRD or DCRD may affect the catalytic activity of PKAC. We examined the effect of purified PCRD_trunc_ and DCRD on PKAC catalytic activity using two different assays with the standard PKA substrate, Kemptide (80). This peptide contains two serines, S^272^ and S^300^, implicated in PKA phosphorylation of Rad that mediates regulation of Ca_V_1.2 (71). Neither PCRD_trunc_ nor DCRD caused any significant changes in PKAC activity (Fig. S6).

### Differences in β1AR modulation of full-length and truncated Ca_V_1.2 channels

As discussed above, in cardiomyocytes the two forms of α_1C_ coexist in an unknown proportion, and it is not known if they are distinctly regulated by PKA. A difference may be expected in light of our discovery of the triple interaction between PKAC and proximal and distal parts of α_1C_ C-terminus. Therefore, we assessed the role of α_1C_-dCT and the proteolytic processing of cardiac α_1C_ using a functional electrophysiological assay. We compared the β1AR regulation of Ca_V_1.2 containing either FL or truncated α_1C_ in a heterologous expression system. To date, *Xenopus* oocyte is the only model cell in which the full adrenergic cascade of regulation of Ca_V_1.2 (starting with the activation of the βAR) has been reconstituted. In a previous study we demonstrated robust, Rad-dependent βAR regulation of both full-length and truncated Ca_V_1.2 by β1AR or β2AR (54). The regulation of Ba^2+^ currents (I_Ba_) by the βAR agonist isoproterenol (Iso) appeared smaller in the case of the FL α_1C_ (54) but in a limited range of current amplitudes, especially for α_1C_-FL. We have therefore decided to characterize the βAR regulation of the two forms of Ca_V_1.2 in more detail and in a wider amplitude range.

Oocytes were injected with RNAs of all channel subunits (α_1C_-FL or α_1C_Δ1821, α2δ1, β_2b_), β1AR and Rad. I_Ba_ was measured every 10 s, by 20-ms voltage steps from a holding potential of -80 mV to +20 mV (Fig. 8A inset), using two-electrode voltage clamp (TEVC). Fig. 8A shows the time course (diary) of a representative experiment. After establishing the voltage clamp, the current initially gradually increased and then often decreased to a stable level, which we considered as the basal I_Ba_ (denoted by the dotted line in Fig. 8A). This process took 10-15 min. Addition of Iso (200 nM) following the stabilization of basal I_Ba_ increased the current; the effect reached its maximum after 5 to 10 min. We collected data from 72 (α_1C_) and 80 (α_1C_Δ1821) cells and, on average, did not observe any statistically significant differences in Iso effect between the full-length and the truncated channel (Fig. 8B). Nevertheless, a closer examination of raw data revealed a negative correlation between the basal I_Ba_ and the Iso effect in Ca_V_1.2 Δ1821 that was absent in full-length channels (Fig. 8C,D). This distinction is further illustrated in Fig. 8E where cells have been subdivided into groups according to basal I_Ba_. Iso caused a significantly higher response in Ca_V_1.2- Δ1821 compared to the FL channel when the basal current was low, but this tendency disappeared and in fact reversed when the current was high (over 700 nA). We have re-analyzed the raw data of the previously reported series of experiments (Fig. 4 in (54)) and found the same general tendencies, with a negative correlation between basal I_Ba_ and fold-increase by Iso in the truncated, but not the FL channel (Fig. S7). Thus, the PKA modulation of FL and Δ1821 Ca_V_1.2 differs mechanistically and involves PCRD/DCRD interaction, as discussed below.

**Figure 8.**
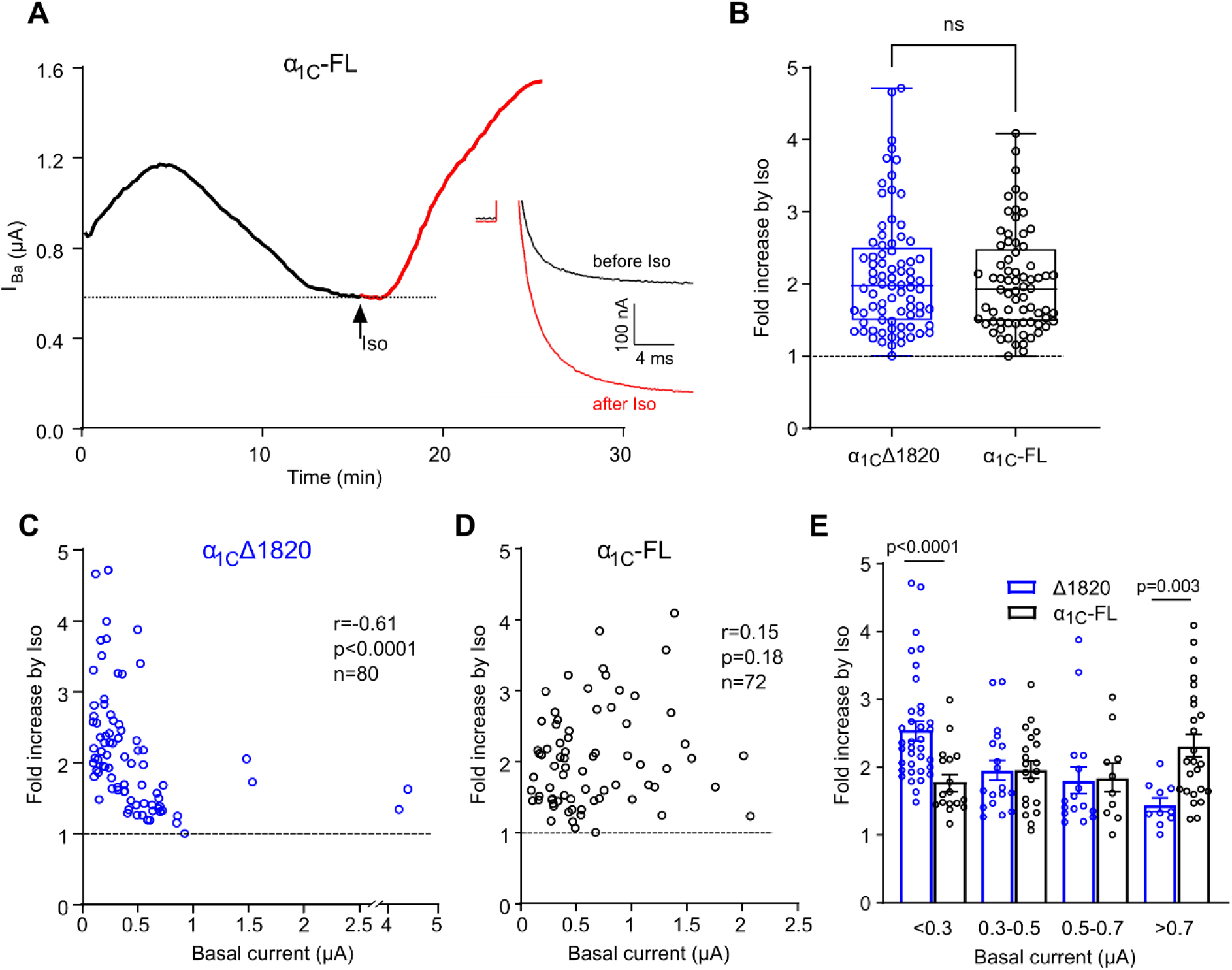
Quantitative differences in β1AR regulation of full-length and truncated Ca_V_1.2. *A*, time course (diary) of changes in I_Ba_ in a representative experiment. Oocytes expressed α_1C_, β_2b_, α2δ1, Rad and β1AR. I_Ba_ was measured by 20-ms depolarization steps from -90 to +20 mV every 10 s, starting shortly after electrode insertion (t=0). Iso (200 nM) was applied at the indicated time (arrow). *Inset* shows current traces before (basal) and at the leak of Iso effect. Most of the capacity transient (an upward deflection preceding the actual inward I_Ba_) has been erased for clarity. *B*, Summary of Iso-induced increase in all oocytes expressing α_1C_Δ1821 (n=80) and α_1C_-FL (n=72). Boxes show 25-75 percentiles and median, whiskers extend to maximal and minimal values. There was no significant difference between the two groups (p=0.59, two-tailed Mann-Whitney test, U=2735). Here and in C, D the dotted line indicates fold increase of 1 (no effect of Iso). *C* and *D*, correlation between basal I_Ba_ and fold increase by Iso in individual cells. Parameters of Spearman correlation analysis for the two distributions are shown in insets. *E*, Comparison of the differences in Iso-induced increase in I_Ba_ of α_1C_Δ1821 and α_1C_-FL for basal I_Ba_ of small, intermediate and high amplitude. In each subgroup of amplitudes, the difference between α_1C_-FL and α_1C_Δ1821 was analyzed by Mann-Whitney unpaired test except for currents >0.7 µA where the data distributions passed the Shapiro-Wilk normality test, and an unpaired t-test was performed.

## Discussion

βAR regulation of Ca_V_1.2 is achieved mainly via phosphorylation of channel-associated Rad (71) by PKAC following its separation from PKAR. Since AKAPs are not essential for this regulation in cardiomyocytes (50,51), we hypothesized that PKA may interact directly with α_1C_. Using several independent methods, we showed here that PKAC interacts directly with α_1C_, and have mapped two regulatory domains on α_1C_-CT, PCRD and DCRD, that bind PKAC in the absence of PKAR. We also show that PCRD and DCRD interact with each other, and addition of DCRD reduces PKAC/PCRD interaction in a dose-dependent manner.

First, in a heterologous expression model (HEK293 cells), we showed co-immunoprecipitation (co-IP) of PKAC-YFP with α_1C_Δ1821, independently of PKAR or dCT expression. Importantly, activation of PKA (by cAMP or indirectly, by forskolin) dramatically reduced co-IP of PKAC with PKAR, as expected. In contrast, the binding of PKAC to α_1C_ did not decrease and even showed a tendency to increase (Fig. 2). These findings suggest a PKAR-independent interaction between α_1C_ and PKAC, although it remains possible that the PKA holoenzyme can also bind α_1C_ in the absence of AKAP. They also offer an explanation to the observation made by a proximity assay, in which PKAC was found in a close association with the channel only following βAR stimulation in cardiomyocytes (53), i.e. after separation of PKAC from PKAR. We propose that PKAC is found in closer proximity to the channel because, when separated from PKAR, it binds to α_1C_-CT.

Next, peptide array analysis of α_1C_ identified segments in α_1C_-CT, specifically within PCRD and DCRD, which interact with PKAC (Fig. 3). The complementary peptide array analysis of PKAC suggested two domains on PKAC that interact with PCRD, one of these domains also interacted with DCRD (Fig. 4). Further studies may be needed to better delineate the exact DCRD and PCRD binding sites in PKAC.

We corroborated the direct interaction of PKAC with PCRD and DCRD using recombinant purified proteins (PCRD_trunc_, DCRD, PKAC) (Fig. 5) and synthetic peptides (Fig. 7) in pull-down experiments, and observed a dose-dependent and saturating binding of PCRD_trunc_ and DCRD to PKAC (Fig. 5). Finally, the interaction between purified PCRD_trunc_ and PKAC was further demonstrated and extended by isothermal titration calorimetry (ITC) (Fig. 6). Although we used truncated PCRD and the exact truncation site is unknown, the pulldown experiments with the PCRD44 and PCRD30 peptides confirmed that the main binding site is located within PCRD_trunc_ and contains the GPEI**RR**AI**S**G motif, where the penultimate serine is S^1700^, known to be phosphorylated by PKA (2). A transient interaction between the PCRD and PKAC could be expected, as it contains the PKAC RRxxS^1700^ target motif (81). The homology of this segment in PCRD to high-affinity PKA-binding pseudosubstrate sites of PKAR and PKI (Fig. S4F) further increase our confidence that this is indeed a major PKA-binding segment in PCRD. In contrast, there are no known or potential PKA phosphorylation sites in the DCRD. Interestingly, we found that both DCRD and PKAC bind to this same PCRD30 peptide.

ITC, which accurately estimates the thermodynamic parameters of protein interactions, determined the K_D_ of PKAC/PCRD_trunc_ interaction in the low-micromolar range (Fig. 6). Interestingly, a recent study did not detect significant binding between PKAC and a short 18-mer peptide RTLHDIGPEIRRAISGDL from PCRD, using ITC (82). This implies that additional residues either upstream or downstream from sequence, present in our PCRD_trunc_ protein, and probably in the PCRD30 peptide, are involved in PKAC binding. Furthermore, there may be an additional binding segment beyond PCRD_trunc_ indicated by the peptide array (Fig. 3), so that the actual affinity of this interaction may be higher than 1 µM. ITC also revealed an apparent stoichiometry of 1.7 that is in line with the presence of two potential PCRD-binding sites in PKAC (Fig. 4). Further study is needed to determine the stoichiometry of PKAC/PCRD interaction. Regardless, our finding that the binding affinity is in the low µM range is in line with dynamic biological regulation (83). Nevertheless, affinity of PKAC for its phosphorylation substrates tends to be lower than micromolar, enabling the fast turnover necessary for efficient catalysis (84). Thus, one to two orders of magnitude lower affinity was found in three best-phosphorylated peptide substrates related to Ca_V_1.2 regulation, one from α_1C_ (containing the S1928 phosphorylation site, by rabbit α_1C_ count) and two from Rad (82). In that study, the 18-mer PCRD peptide was a poor substrate for phosphorylation by PKAC. Therefore, the functional role of the PCRD-PKAC interaction may indeed not be phosphorylation of S1700. Rather, we posit that although PKAC will not be obligately “anchored” to the PCRD site, it may be enriched in that nanodomain due to kinetic scaffolding (85). This potential mechanism facilitates localized signaling by restraining PKAC diffusion (86).

The binding of PKAC to both PCRD and DCRD is intriguing, since the interaction between the two domains is envisaged as part of the mechanism by which α_1C_-dCT inhibits channel’s gating. A direct interaction between PCRD and DCRD was deduced from a structure-function study and predicted, using a computational model, to exist via salt bridges between specific residues (including R^1696^ and R^1697^ of PCRD) on both domains (24). Our results (Fig. 7B) present the first experimental evidence for such a direct interaction. In support, Lyu et al. (87) demonstrated interaction between two large segments of the CT, one including the EF, pre-IQ, IQ and PCRD domains in the proximal CT and the other including DCRD and CCD (downstream of DCRD) in the dCT. This interaction was inhibited by CaM in a Ca^2+^-dependent manner. In addition, the authors reported an interaction between segments of proximal and distal CT that contained neither PCRD nor DCRD (87). In Ca_V_1.3 channels three domains were suggested to interact to compete with CaM: PCRD, DCRD and IQ_V_ (preIQ_3_-IQ domain), modulating both activation and inactivation of the channel (88). Finally, PCRD/DCRD interaction was proposed to participate in Ca_V_1.2 regulation by PKA through the phosphorylation of S^1700^ (2), possibly in concert with the Rad-dependent regulation (72,73), though the role of phosphorylation of S^1700^ remains debated (25,53,89). Overall, it appears that PCRD/DCRD-dependent and additional interactions between the proximal and distal CT in α_1C_ and α_1D_ (Ca_V_1.3) profoundly regulate the channel’s function and possibly its regulation by Ca/calmodulin and PKA.

It was even more intriguing to find that the same 30-a.a. (and possibly shorter) motif within PCRD interacts both with PKA and DCRD (Fig. 7), and that DCRD and PCRD share a common binding site on PKAC (Fig. 4). This suggested a compound interaction module in which all three binding partners compete for common binding sites, as supported by the triple interaction experiment that showed a concentration-dependent reduction in PKAC/PCRD30 binding by DCRD (Fig. 7). We thus propose that PCRD serves as a “hub” for these interactions, and that the dissociated PKAC monomer remains a part of Ca_V_1.2 multimeric complex.

The tripartite interaction of the two important regulatory C-terminal domains of α_1C_ with PKAC indicated a potential physiological role, in which the dCT may modulate the action or abundance of PKAC. We ruled out the scenario in which binding of PCRD or DCRD regulates the catalytic function of PKAC (Fig. S6). We then addressed the role of dCT by studying the regulation of the channel by β1AR activation in *Xenopus* oocytes, the only heterologous expression system in which the entire cascade can be reconstituted (54). We expressed either Ca_V_1.2-FL or Ca_V_1.2Δ1821 along with β1AR and Rad. On average, Ca_V_1.2-FL and Ca_V_1.2Δ1821 were similarly upregulated by β1AR activation by the agonist, isoproterenol (Iso). However, an in-depth analysis of the regulation according to basal current before Iso application (which may correspond to the number of functional channels) revealed a difference between the two forms of the channel. We found a negative correlation between the basal current of Ca_V_1.2Δ1821 and the extent of Iso-induced regulation. No correlation was found in oocytes expressing Ca_V_1.2-FL (Fig. 8). This implies a specific function of the dCT in βAR regulation.

This analysis has several limitations, such as possible differences in the ratio of expressed channel subunits to Rad (which may affect the extent of βAR regulation), yet the reproducibility of these results in two separate series of experiments (Fig. 8 and Fig. S7) supports the validity of our observations and highlights the differences in the physiological regulation of the two forms of the channel by PKA.

Our observations are in line with a previously reported similar negative correlation between the Ca_V_1.2 basal current and the magnitude of βAR regulation in cardiomyocytes (90). The negative correlation was observed both in untreated cardiomyocytes and in cells overexpressing Ca_V_β subunits. The authors speculated that there may be a limiting factor, such as a final number of permissive sites on the membrane, where PKA-mediated upregulation of Ca_V_1.2 can occur. When the amount of channel protein exceeds the amount of the limiting factor, a further increase in the channel’s surface density results in a subpopulation of Ca_V_1.2 irresponsive to PKA. Among such factors, we can think about available PKA or Rad; indeed, not all channels are constitutively associated with Rad (62,70). We further propose that the existence of the triple PCRD/DCRD/PKAC complex may also stabilize the PKAC – Ca_V_1.2 association in the full-length α_1C_. The secondary Ca_V_β-binding domain in the dCT, located very close to DCRD (Fig. 1), may further contribute to the formation of the macromolecular complex by physically or “kinetically” scaffolding both Ca_V_β and the associated Rad in the proximity of α_1C_-CT and PKAC. In the truncated form of the channel, Ca_V_1.2Δ1821 such multimolecular complex may be absent or less tight, thus either Rad or PKAC may not be sufficiently recruited to channel’s vicinity, becoming limiting factors.

As noted before, a large subpopulation of cardiac Ca_V_1.2 is truncated at the dCT, but the extent of truncation is uncertain. Since a negative correlation between the basal current and Iso effect was observed in native guinea pig and rat cardiomyocytes, our observations support the notion that, in these species, most of Ca_V_1.2 is actually truncated.

In summary, we show that PKAC binds two regulatory domains on the CT of α_1C_ (PCRD and DCRD) independently of PKAR or the need for AKAPs. We also show that these two domains interact with each other. Finally, we propose that dCT coordinates the function of the multimolecular complex involved in βAR regulation of Ca_V_1.2, by supporting the scaffolding of the macromolecular complex of Ca_V_1.2 with PKAC and Rad, which is essential for an efficient βAR regulation of the channel. In the dCT-truncated channel the efficiency of PKA regulation is at least as high as in the FL channel but depends on limiting factor(s).

## Materials and methods

### Animal use and ethical approval

Experiments were approved by Tel Aviv University Institutional Animal Care and Use Committee (IACUC permit # 01-20-083). Female frogs were kept in tanks filled with dechlorinated water that was exchanged 3 times a week, at 20±2 °C on 10 h light/14 h dark cycle. Frogs were anaesthetized in 0.17-0.2% tricaine methanesulfonate (MS-222), and portions of ovary were removed through an incision on the abdomen. The incision was sutured, and the animal was held in a separate tank until it had fully recovered from the anesthesia, and afterwards was returned to a separate tank for post-operational animals. The animals did not show any signs of post-operational distress and were allowed to recover for at least 3 months until the next surgery. Following the final collection of oocytes (or the 4^th^ surgery as defined by the IACUC rules), anaesthetized frogs were sacrificed by decapitation and double pithing.

### DNA constructs and RNA

For transient expression in HEK293T cells, the following constructs were used: mouse PKAC-YFP in pcDNA3 (mouse PKAC, GenBank: NM_008854, fused to eYFP, GenBank: GQ221700.1) PKA-RIIβ-HA (GenBank: NM 001020020.1; the HA tag was fused in frame at the end of the C-terminu). For protein purification from *E. coli*, the following constructs were used: His-tagged Cα-subunit of PKA (His-PKAC, GenBank:NM 008854.5), glutathion-S-transferase (GST)-fused PKA-RIIβ (GST-PKAR; GenBank: NM 001030020) (76), GST-fused Cα-subunit of mouse PKA, GST-PKAC (GenBank: NM008854.4) (kind gift from S.S. Taylor). In addition, we have produced DNA constructs for protein production in *E. coli* in pET-Duet vector, encoding the following proteins: 1) His_8_-SUMO-PCRD, with a 3 a.a. segment MGS followed by an 8-His (His_8_) tag fused to SUMO (a.a. 1-81 of human Chain B of small ubiquitin-related modifier 2; PDB: 2IO3_B) further fused via a ENLYFQG liker (a TEV protease cleavage site) to PCRD (a.a. G1671-T1751 of rabbit α_1C_). 2) His_8_-DCRD-Myc, with an MGS-His_8_ tag fused via a TEV digestion site to DCRD (a.a. R2058-S2120 of rabbit α_1C_) followed by a Myc tag, EQKLISEEDL.

RNA for *Xenopus* oocyte injections was transcribed *in vitro* using previously described protocols and DNA constructs (54,76): human Rad (NP_001122322), cardiac long-N terminus isoform of rabbit α_1C_ (GenBank: X15539), rabbit Ca_V_β_2b_ (GenBank: X64297.1), rabbit α2δ1 (GenBank: M21948), mouse β1 adrenergic receptor (NP_031445.2). For oocyte expression, the coding sequences of these DNAs were inserted into pGEM-SB or pGEM-HJ vectors containing the 5’ and 3’ untranslated regions of *Xenopus* β-globin (91).

Ca_V_1.2 was expressed in *Xenopus* oocytes in full subunit composition α_1C_+α2δ+β_2b_, by injection of equal amounts, by weight, of RNAs of each subunit. To obtain similar ranges of amplitudes of I_Ba_ for α_1C_-FL and α_1C_Δ1821, and to ensure optimal Rad-dependent regulation by Iso, we tested a range of RNA doses for the channel subunits and Rad. For experiments shown here we chose the following combination of RNA doses, per oocyte: α_1C_: 2 ng for α_1C_-FL and 1 ng for α_1C_Δ1821; 0.5 ng Rad RNA for α_1C_-FL and 1 ng Rad for α_1C_Δ1821. RNA of β1AR was 10-20 pg/oocyte.

### Electrophysiology in Xenopus oocytes

Oocytes were defolliculated by collagenase treatment (92), injected with the desired combinations of RNAs, and incubated for 3 days at 20–22 °C in NDE solution (in mM: 96 NaCl, 2 KCl, 1 MgCl_2_, 1 CaCl_2_, 5 Hepes, 2.5 pyruvic acid, and 0.1 gentamycin sulfate). Recordings started 3 days after RNA injection and conducted over the next 2 days. Comparison of basal currents between test groups were performed on recordings from the same day. Results of “before-after Iso” analysis of each oocyte (Iso) were pooled from all the recordings performed over 2-3 days. Oocytes of the various experimental groups were tested intermittently over the entire duration of the experiment to normalize variability, as protein levels increase over time. Oocytes were placed in a recoding chamber (solution volume ∼100 µl) perfused at a steady rate, that did not change throughout the experiment, with 40 mM Ba^2+^ solution (in mM: 40 Ba(OH)_2_, 50 NaOH, 2 KOH, and 5 Hepes, titrated to pH 7.5 with methanesulfonic acid). The chamber was connected via 3M KCl bridges and silver chloride-coated electrodes to the ground inputs of the virtual ground module of the Geneclamp 500 amplifier (Molecular Devices, Sunnyvale, CA, USA). Currents were recorded using two-electrode voltage clamp in virtual ground mode (93), digitized with Digidata 1440A, and acquired and analyzed with the pCLAMP software (Molecular Devices, Sunnyvale, CA, USA). I_Ba_ was elicited by depolarizing pulses from a holding potential of -80 mV to +20 mV applied at 10 s intervals. The voltage step to +20 mV was preceded by a hyperpolarizing prepulse to -90 mV (100 ms), to estimate the leak conductance. Net I_Ba_ was calculated by subtracting the calculated leak currents during the analysis session. Iso (Sigma-Aldrich, I5627) was diluted in water, aliqoted and stored in 100 mM stock solution at -20 °C. Perfusion of isoproterenol (200 nM) began after attaining a stable peak current for about 2 minutes (see Fig. 8).

### Generation and culture of the inducible mouse Ca_v_1.2a HEK293 cell line

To generate the inducible mouse Ca_v_1.2a 293 cells, stop codon was introduced after Val1821 in mouse Ca_V_1.2a (isoform 7, NM_001255999.2) and its cDNA subcloned into the pcDNA5/FRT vector. After transfection the Flp-In TRex HEK293 (Flp-In-293, Invitrogen) cells were selected in the presence of Hygromycin B (100 µg/ml) and single clones were grown and analysed by Western blot before and after tetracycline-induced mouse Ca_V_1.2a cDNA expression (Fig. S1A). Cells were induced for 18 h with 1 μg /ml tetracycline (Invitrogen). The cells were cultured in Dulbecco’s Modified Eagle Medium (DMEM 41965-039, Gibco, Thermo Fisher Scientific, Waltham, USA), supplemented with 10% fetal bovine serum (FBS, Gibco), and 200 mM L-Glutamine, 100 µg/ml Hygromycin B and 100 U / ml Penicillin with 100 μg/ ml Streptomycin at 37 °C and 5% CO_2_.

HEK293 cells were maintained in Dulbecco’s Modified Eagle’s Medium (DMEM) supplemented with 10% Fetal Bovine Serum (FBS), 100 U/ml penicillin, 10 mg/ml streptomycin and 2 mM L-Glutamine (Biological Industries) at 37 °C with 5% CO_2_. Transient transfection with the calcium-phosphate was 48 h before the cells were collected.

### Immunoprecipitation from HEK293 cells

For activation of adenylyl cyclase, cells were incubated for 10 minutes with DMSO (1:1000) or with 25 μM forskolin (Alomone, stock 10 mM in DMSO). Cells were scrapped in PBS + 0.5 mM phenylmethylsulfonyl fluoride (PMSF) at 4 °C and cell pellets were stored at -80 °C. Cell pellets were solubilized in *lysis buffer*: 1 % Triton x100, 20 mM Tris pH 7.5, 20 mM EDTA, 10 mM EGTA + protease inhibitors (cOmplete, Roche), 25 mM β-glycerol-phosphate, 2 mM orthovanadate and 0.5 mM PMSF, for 30 min on ice then cleared by centrifugation at max speed. Supernatants were transferred for protein quantification (Lowry assay) and immunoprecipitation (IP). For IP, 900 µg total protein were dilutes in 300 µl lysis buffer, and 25 µg total protein were taken as input. Samples were incubated with 2 µl IP antibody at 4 °C in a rotating device, over-night. Protein A-Sepharose beads (GE17-0780-01, 50 µl) were taken for each tested sample from the slurry and incubated overnight with 0.5 % BSA in PBS, then washed with PBS and lysis buffer. Then an equal amount of washed beads was divided per sample and incubated at 4 °C for 90 min in a rotating device. Beads were washed on a rotating device, at RT, three times with 1 ml *washing buffer*: 1 % Triton x100, 20 mM Tris pH 7.5, 150 mM NaCl and 0.5 mM PMSF, and a final wash with 20 mM Tris pH 7.5 and 0.5 mM PMSF. When indicated, 1 mM cAMP (Sigma-Aldrich, stock 20 mM in DDW) were added to the washing buffer. Proteins were eluted by incubation with sample buffer for 5 min at 65 °C, centrifuged and the clear supernatant was taken for acrylamide-gel loading and Western blot analysis.

### Electrophysiology in HEK293 cells

Currents were recorded using whole-cell patch-clamp configuration, and sampled at 10 kHz. Signals were amplified using an Axopatch 200B patch-clamp amplifier, digitized with Digidata 1440A and analyzed using pCLAMP 10 software (Axon Instruments). External solution contained (mM): 150 Tris, 10 glucose, 1 MgCl_2_ and 10 BaCl_2_ (adjusted to pH 7.4 with methanesulphonic acid). The intracellular solution contained (mM): 135 CsCl_2_, 10 EGTA, 1 MgCl_2_, 4 MgATP, 10 Hepes (pH 7.3, adjusted with CsOH).

### Production and purification of recombinant proteins

Recombinant proteins were all expressed in *E. coli* (BL21-DE3 and derivatives and produced using standard methods, essentially as described (76,94). Freshly transformed colonies were inoculated into 2xYT media, grown at 37° C, induced with IPTG (0.25-1 mM) at OD=0.4- 0.6. Growth after induction was at 16 – 24 °C overnight, or, for GST-PKAC, 6.5 hours at 24 °C. Bacterial cells were harvested by centrifugation, frozen and stored at -20 or -80° C for later use.

For His-tag proteins, defrosted bacterial paste was resuspended in lysis buffer: potassium phosphate, pH 8 (50 mM), 0.1 M NaCl, β-mercaptoethanol 5 mM or PBS pH 7.8 with β- mercaptoethanol (2 mM). DNase, lysozyme, and PMSF and protease inhibitor cocktail (solution or tablets) were added to facilitate lysis and inhibit proteolysis. Cells were lysed either by microfluidizer or sonication. The lysate was then centrifuged at high RCF for 30 -60 min and soluble material was loaded on a Ni-NTA metal-chelate column. The resin was then spun down at 3000 rpm and the supernatant removed. The column was washed with buffer A (lysis buffer including 10 mM imidazole) and HisTag protein was eluted with buffer B (lysis buffer including either 250 or 150 mM imidazole). Fractions were pooled, concentrated and loaded onto a SEC column (Superdex 75) equilibrated with either potassium phosphate pH 7.5 (20 mM), KCl (20 mM), DTT (2 mM) or PBS pH 7.8 with β-mercaptoethanol (2 mM).

GST-fused proteins were purified as described (76). Lysis was as above. Soluble lysate was then loaded on a glutathione-agarose column and eluted with glutathione. The eluate was then pooled, concentrated and further purified using size-exclusion chromatography on Superdex-75 in the following buffer, in mM: 20 KH2PO4, 20 KCl, 2 DTT, pH 7.5. After purification, protein fractions were concentrated, frozen by flash freezing in liquid nitrogen and stored at -80 °C for later use.

### Reversed-phase (RP) HPLC

Reversed-phase (RP) HPLC of uncleaved and cleaved forms of His-SUMO PCRD was performed on an JASCO PU-1580i HPLC system equipped with a UV detector (PU-1575) and flow cell (JASCO Inc. Japan). The mobile phase included water with 0.1% trifluoroacetic acid (TFA) in solvent A and 100% acetonitrile (ACN) with 0.1% TFA in solvent B. Vydac C4 semi-preparative RP column with 5 μm particle size and column dimensions 240 mm × 10 mm were used for the HPLC runs. The columns were operated at room temperature with a flow rate of 0.5 mL/min. The column eluate was analyzed by UV detector and collected manually and further run on 12% SDS-PAGE gels for analyses. Each HPLC run lasted for approximately 80 min with a 0-100% gradient of Water:ACN was achieved in the end of 80 minutes.

### Nano LC ESI-MS/MS mass spectrometry and raw data analysis

Lyophilized protein was reconstituted in destilled water at a concentration of 12.5 µg/µl. 15 µg were digested in solution with SMART Digest™ Trypsin magnetic beads (Thermo Fisher) at a total volume of 200 µl at 70 °C for 2 h. 10 µg of reconstituted protein was digested in 0.05 N HCl with 1 µg pepsin (Promega) without prior reduction/alkylation at a volume of 200 µl 50 °C for 3 h. Nano-LC-ESI-MS/MS was performed as described previously (95) using a LTQ Orbitrap Velos Pro coupled to Ultimate 3000 RSLC nano system equipped with an Ultimate3000 RS autosampler, (ThermoFisher Scientific, Dreieich, Germany). Peptides were separated on a reversed phase column (nano viper Acclaim PepMap capillary column, C18; 2 µm; 75 µm × 50 cm, ThermoFisher) at a flow rate of 200 nL/min increasing organic acetonitrile phase for 60 min as described before. We used a combination of alternate the collision induced dissociation (CID) fragmentation and high energy collision dissociation (HCD) top3 method, where full scan MS spectra (m/z 300–1700) were recorded in the orbitrap analyzer with resolution of r = 60,000. The 3 most intense peptide ions with charge states >2 were sequentially isolated and fragmented in the HCD collision cell with normalized collision energy of 30%. The resulting fragments were detected in the Orbitrap system with resolution r = 7,500. For the collision induced dissociation (CID) MS/MS top3 method full scan MS spectra (m/z 300–1700) were acquired in the Orbitrap analyzer using a target value of 10^6^. The 3 most intense peptide ions with charge states >2 were fragmented in the high-pressure linear ion trap by low-energy CID with normalized collision energy of 35%.

Tryptic and peptic peptides were analyzed by Mascot/Proteome Discoverer 1.4 (Thermo Scientific). Raw files were searched against a SwissProt database containing 16.992 *mus musculus* entries and the manually added sequence of tagged PCRD_trunc_. MS^2^ spectra were matched with a mass tolerance of 7 ppm for precursor masses and 0.5 Da for peptide fragment ions. Database search was done with “no specific” enzyme digestion. Dynamic peptide modifications as oxidation at methionine, deamidation at asparagine and glutamine and acetylation, methylation and dimethylation at the N-terminus were allowed.

### Pull-down of purified proteins and synthetic peptides

Procedures were essentially as described (76). Interaction of His-fused proteins with other purified proteins (e.g. GST-PKAC) or synthetic peptides was studied by pulldown using Ni-NTA affinity resin (HisPur, Thermo scientific # 88221). GST-fused proteins were pulled down on glutathione-Sepharose beads (Amersham Biosciences). The binding buffer contained 150 mM KCl, 5 mM MgCl_2_, 50 mM Tris-HCl (pH=7.5), 0.025% TWEEN-20 or 0.5% CHAPS, and 1 mM EDTA. The binding reaction (300 µl total volume) was initiated by mixing the interacting proteins or peptides. After 60 min incubation on a shaker at room temperature (or 2 hours at 4°C when pull-down was done with GST-fused proteins), 5 µl of the reaction mixture was removed to later visualize the loaded proteins ("input"). At this stage, for pull-down on Ni-NTA beads, 3 µl of 1 M imidazole and 30 µl of Ni-NTA agarose beads (Thermo Scientific, MA, USA) were added to the mixture and incubated for 30 min at 4-8°C, then washed 3 times with 500 µl of the incubation buffer containing 10 mM imidazole. Elution was done by 10 min incubation at room temperature with 30 µl of the incubation buffer containing 250 mM imidazole. For pull-down on glutathion beads, GST fusion proteins 30 µl were immobilized on glutathione-Sepharose beads (30 µl; Amersham Biosciences) for 60 min at 4 °C, washed three times with 500 µl of the binding buffer. Then the buffer was removed, 20 µl of ×2 sample buffer (2% SDS, 10% β-mercaeptoethanol, 8% glycerol, 50 mM Tris, pH 6.5) were added to the beads, followed by incubation for 5 min at 95°C to elute the bound protein from the beads, addition of 20 µl water, and centrifugation for 2 min at 1000 rpm in a tabletop centrifuge. The protein eluates from Ni-NTA or glutathion beads and the “input” were separately subjected to SDS-polyacrylamide (12%) gel electrophoresis (SDS-PAGE). Running buffer contained: 25 mM Tris, 192 mM glycine, 1% SDS, pH 8.3. The synthetic fluorescein-labeled peptides were pulled down in the same way, but dissolved in SDS-free sample buffer, and instead of standard SDS-PAGE we used 0.75 mm thick, 16.5% Tricine gels. The running buffers contained: cathode buffer 10X (upper buffer): 0.1 M Tris, 0.1 M Tricine, 0.1% SDS, pH 8.25; anode buffer: 0.2 M Tris, pH 8.9. The gels were imaged immediately after the end of electrophoresis with Fusion FX7 (Witec AG, Sursee, Switzerland) (excitation 500-550 nm, emission filter F595Y3) and subsequently stained with Coomassie Briliant Blue G-250 (Bio-Rad #1610406) 0.1%, 8% ammonium sulfate, 1% orthophosphoric acid, 20% ethanol.

### Western blot and antibodies

Nitrocellulose membranes were blotted with protein samples and incubated for 1 h in 5 % skin milk in PBS, supplemented with 0.5 % Tween 20. Incubation with the primary antibody was overnight at 4 °C, and for secondary antibody, 1 h at room temperature. Membranes were incubated with chemiluminescent HRP substrate solution (SuperSignal West Pico, Thermo Scientific) and exposed to a film. The following antibodies were used as indicated: GFP, G1544 Sigma Aldrich (rabbit); Ca_V_1.2 (α_1C_), ACC003, Alomone; PKAC SC-903 (PKAα), Santa Cruz (rabbit); Na_V_1.1, ASC-001 Alomone; HA antibody, Santa Cruz SC7392, mouse; His6 Peroxidase, Roche 11965085001; Cavβ (in-house for β3 and β2; PMID: 21357697). Secondary antibodies: goat anti rabbit (Jackson, 111-035-1440); mouse anti rabbit light chain (Jackson, 211-032-171). Densitometry of bands was using ImageJ (NIH). Normalized level of co-IP was the co-IP band intensity divided by the band intensity of IP, e.g. when PKAC-YFP was immunoprecipitated, co-IP normalized level was calculated as: α_1C_= (α_1C_/PKAC-YFP); (PKAR-HA)/(PKAC-YFP). Normalized co-IP level for a treatment was divided by normalized co-IP of control (e.g. DMSO).

### Peptide spot array of α_1C_ cytosolic segments and of PKAC

Peptide array of N-terminal and C-terminal parts of rabbit α_1C_ (X15539) and an N-terminal segment of human α_1C_ (XP_006719080.1) was spot-synthesized as 25-mer peptides overlapping sequences, shifted by 5 a.a. along the sequence, using AutoSpot Robot ASS 222 (Intavis Bioanalytical Instruments, Cologne, Germany), as described previously (96). Peptides were generated by automatic SPOT synthesis and blotted on a Whatman membrane. The interaction with spot-synthesized peptides was investigated by an overlay assay. Following blocking (1 h, RT) with 5% BSA in 20 mM Tris and 150 mM NaCl with 0.1% Tween-20 (TBST), 0.1-0.02 µM purified His-PKAC were incubated with the immobilized peptide-dots, over-night at 4 °C. PKAC was detected by PKAC antibody (sc-903) and HRP-coupled secondary antibody incubated with 5% BSA/TBST, and the membrane was exposed to a sensitive film or imaged with Fusion FX7, as for immunoblotting. Mouse PKACα (NM_008854) was spot-synthesized as above for α_1C_ and incubated with 0.1 µM recombinant His-DCRD-Myc or His-SUMO-PCRD at 4 °C overnight in 50 mM Tris pH 7.4, 5 mM MgCl_2_. Successful binding was analyzed using anti-His Tag antibody (PCRD, Invitrogen #MA1-135) or anti-Myc Tag (DCRD, Cell Signaling Technologies #2276).

### Isothermal Titration Calorimetry

Isothermal titration calorimetry was measured on a MicroCal PEAQ-ITC (Malvern) to examine the interaction between HisTag-SUMO PCRD and HisTag-PKAC. HisTag-SUMO-PCRD and HisTag-PKA were both prepared in the same buffer (PBS pH 7.8 supplemented with 2 mM β-ME) and were degassed before each experiment. Titrations was performed by injecting of His-SUMO-PCRD [400 μM] (syringe) into of His-PKAC [20 μM] (cell) at 25 °C. Each experiment consisted of 13 injections and the time interval between each injection was 150 seconds. The cell volume was ∼200 microliter while the syringe contained ∼75 microliter prior to each experiment. Three control experiments were performed under the same conditions and were subtracted from the experimental dataset: *(i)* HisTag-SUMO-PCRD [400 μM] in syringe and buffer in cell; *(ii)* buffer in syringe and HisTag-PKAC [20 μM] in cell; *(iii)* buffer versus buffer. Data obtained were fit to a one site binding model with the manufacturer’s software.

### Cook Assay

Human PKA-Cα was overexpressed in E. coli BL21(DE3) cells after induction with 0.4 mM isopropyl-β-d-thiogalactoside (IPTG) for 16 h at room temperature using the expression vector pRSETb-hPKACα and then purified by affinity chromatography using an IP20-resin as described earlier (97). Kinase activity of human PKAC in the presence of the two channel fragments was determined in a coupled spectrophotometric assay as described previously (98). The assay buffer contained 100 mM MOPS (pH 7), 10 mM MgCl_2_, 1 mM phosphoenolpyruvate, 1 mM ATP, 2.5 mM β- Mercaptoethanol, 15 U/mL lactate dehydrogenase (Roche Diagnostics GmbH, Mannheim, Germany), 8.4 U/mL pyruvate kinase (Roche Diagnostics GmbH, Mannheim, Germany), 0.1 mg/mL BSA and 200 mM NADH. For the measurement, 1 µM or 10 µM of His-DCRD-Myc or His-SUMO-PCRD were added to 100 µL assay mix. 30 nM of human PKAC was added, and the reaction started by addition of 260 µM S-Kemptide (LRRASLG; GeneCust) at room temperature. NADH depletion was measured using a spectrophotometer (Specord 205, Analytic Jena) at a wavelength of 340 nm for 30 sec. The slope of each measurement was calculated in the WinAspect software (Informer Technologies) and the values exported to Graphpad Prism 9.0 for statistical analysis.

#### ADP-Glo Assay

The ADP-Glo^TM^ Assay (Promega, Germany) is a luminescence-based kinase assay in which the amount of produced ADP is proportional to the luminescence signal. Here, 20 nM bovine PKAC was incubated with Kemptide as substrate in PKA kinase buffer (40 mM Tris-HCl, pH 7.4, 20 mM MgCl_2_, 0.1 mg/mL BSA, 50 µM DTT). 10 µM of His-DCRD-Myc or His-SUMO-PCRD were added to the reaction mix. To inhibit kinase activity 30 µM of H89 were added. Samples were diluted in PKA kinase buffer and reactions set up in a 384 well plate. Kinase reaction was done for 5 minutes at room temperature. The reaction was stopped by addition of 5 µL ADP Glo reagent and the plate incubated for 40 min at room temperature. 10 µL of Kinase detection reagent was added and the plate incubated for 30 min at room temperature. The luminescence was recorded using Tecan plate reader (Tecan, Mannedorf, Switzerland) with an integration time of 0.5 sec. Antibodies used: mouse anti-Hsp90 (Enzo Life Science, #SPA-830); mouse anti-GFP Tag (Thermo Fisher Scientific, #MA5-15256); rabbit anti-pPKA substrate (Cell Signaling Technology, #9621).

#### Statistical analysis

Statistical analysis was performed in GraphPad Prism (GraphPad, La Jolla, Ca, USA). Data were tested for normality of distribution using the Shapiro-Wilk test. In case of normal distribution, pairwise comparison was done by t-test and multiple comparisons by one-way ANOVA with post hoc Bonferroni or Sidak-Holm correction. If the data did not pass normal distribution test, they were analyzed using Mann-Whitney (pairwise) and Kruskal-Wallis non-parametric ANOVA tests.

## Supporting information

***Figure S1, related to Fig. 2.*** Generation of HEK293 cells stably transfected with α_1C_ truncated at 1821 and Ba currents in these cells.

***Figure S2, related to Fig. 3.*** The full peptide array membrane of α_1C_ N-and C-terminal 25 a.a. peptides.

***Figure S3, related to Fig. 4.*** His-DCRD-Myc: purification from E. coli.

***Figure S4, related to Fig. 4.*** His-SUMO-PCRD: purification and analysis.

***Figure S5, related to Fig. 5***. SUMO does not bind PKAC.

***Figure S6***. PCRD and DCRD do not alter the catalytic activity of PKAC.

***Figure S7, related to Fig. 8.*** Quantitative differences in β1AR regulation of full-length and truncated Ca_V_1.2: reanalyzing the published results.

## Acknowledgments

We that Alomone Labs (Jerusalem) for the gift of several antibodies.

## Funding

This study was supported by the following research grants:

German-Israeli Foundation for Scientific Research & Development (GIF) grant I-1452-203.13/2018 (ND, VF, EK, SW, JAH)

Deutsche Forschungsgemeinschaft (DFG): KL1415/14-1 (EK)

DFG, FE 629/2-1 (CFT)

DFG, CRC 894 (VF)

DFG, FL 153/10-2 (VF).

Program-project grant, TP A06, 394046635 – CRC 1365 (EK)

## Author contribution

SO, MK, VF, JAH, EK, SW, ND, FWH conceived the study and supervised experiments; SO, TS, MK, TP, SS, VT, DTT, GS, LV, CFT, TKR, DB planned, performed and analyzed experiments; SW, ND, JAH, EK, SO wrote the paper. All authors read and participated in the editing of the manuscript.

## Data and materials availability

All data are presented in the paper. Materials (new DNA constructs) will be available upon request.

## Supporting information

**Figure S1, related to Fig. 2.**
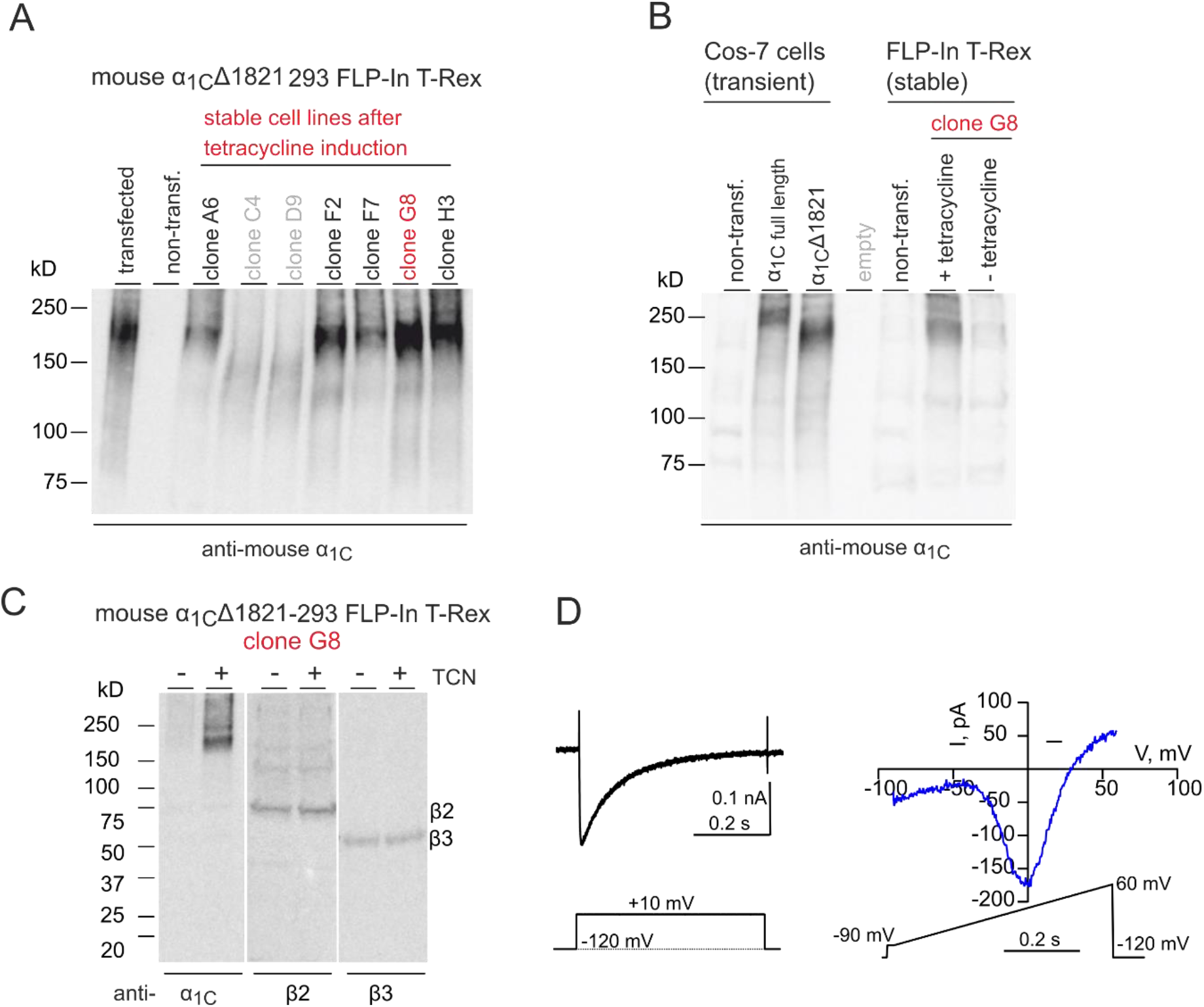
Generation of HEK293 cells stably transfected with α_1C_ truncated at 1821 and Ba currents in these cells. *A,* Seven independent stable cell lines were generated, and tetracyclin-induced α_1C_Δ1821 was detected by Western blot. Clone G8 had the highest α_1C_Δ1821 expression and was selected for all following experiments. B, Western blot analysis of full length α_1C_ vs. α_1C_Δ1821 in Cos-7 cells transiently expressing α_1C_ and in the G8 cell line stably expressing α_1C_Δ1821 (only when induced by tetracycline). Note that the tetracycline-inducible cells contain endogenous Ca_V_β_2_ and Ca_V_β_3_, which probably contribute to the formation of the functional channel. *C,* Exemplary traces from tetracycline-induced G8 clone of typical Ba^2+^ current waveform and current-voltage relationship. Left, currents were elicited by a voltage step from resting potential of -120 mV to 10 mV. Right, voltage ramp protocol from -90 to 60 mV showed a peak current at ∼ 0 mV.

**Figure S2, related to Fig. 3.**
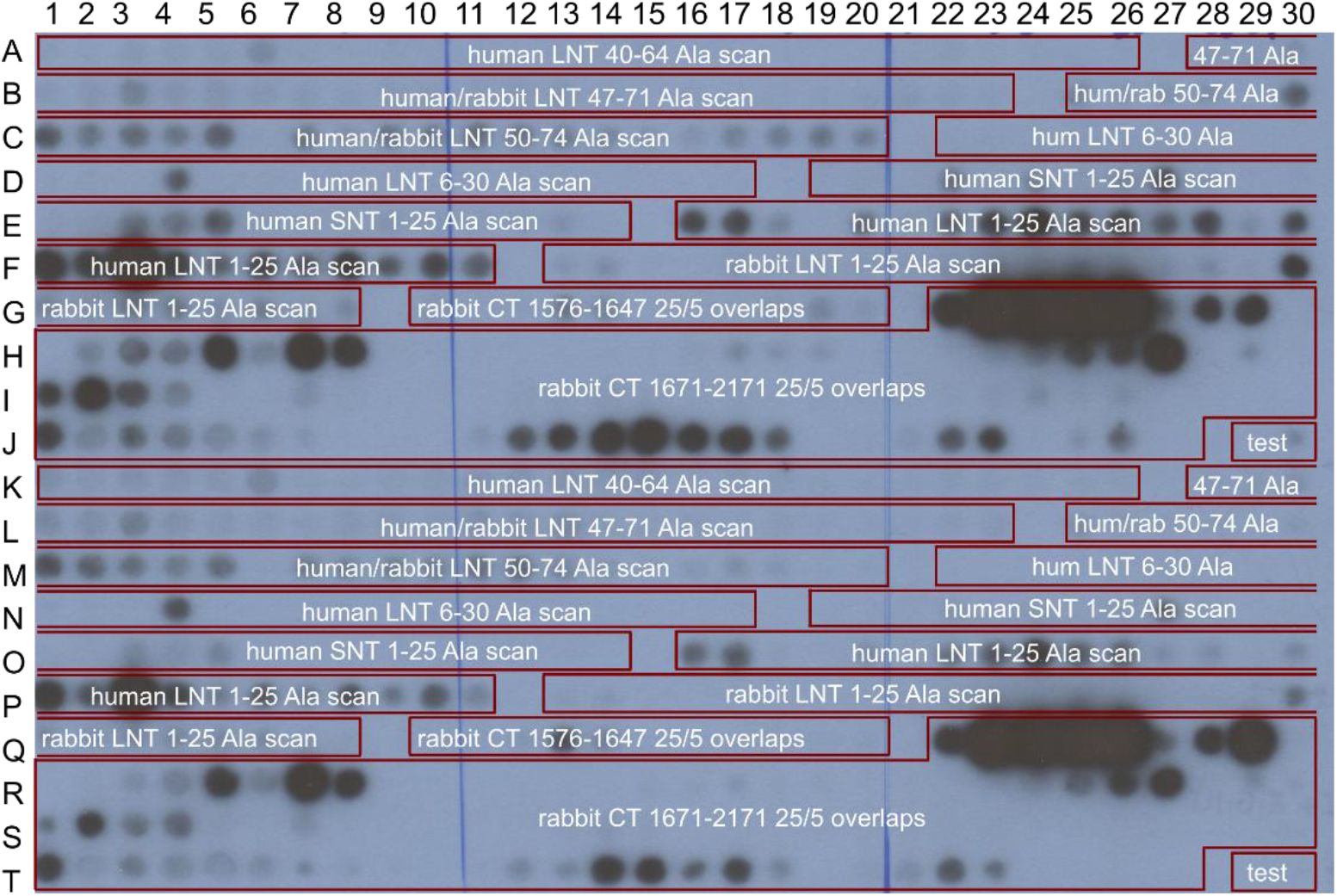
The full peptide array membrane of α_1C_ N-and C-terminal 25 a.a. peptides. Lines K-T present a duplicate of the array of lines A-J. The image shows the result of the overlay with 0.1 µM PKAC (lines A to J) or 0.02 µM PKAC (lines K-T), followed by PKA detection with PKAC antibody (1:1000) and secondary anti-rabbit antibody (1:40000). Each of the 25 spot-long framed stretches in A1-G8 shows alanine scans of the indicated NT peptides. LNT, long-NT isoform; SNT, short-NT isoform. The notation human/rabbit corresponds to stretches of 100% homology between human and rabbit α_1C_. In each stretch, the leftmost spot is the WT peptide and the following spots corresponds to alanine mutations of consequent a.a. (e.g. spot A2 is Ala mutant of the first a.a., spot A3 of the second a.a., and so on). Note that, although several strongly labeled spots are observed within the stretch corresponding to the human a.a. 1-25 peptide, the strongest spots correspond to some of the Ala mutations, and the WT peptide (E16) shows only a modest labeling. Moreover, the overlapping peptide a.a. 6-30, and the highly homologous rabbit 1-25 peptide, show no labeling. The part of the array after alanine scans, G10-J27, comprises 25- mer peptides with 20 a.a. overlaps (5 a.a. shift between peptides) corresponding to WT rabbit α_1C_ CT sequences. The major part of the CT, a.a. 1576-2171 (excluding the IQ domain, a.a. 1648-1670), is the same as shown in Fig. 2A. Note no labeling in the proximal CT, a.a. 1576-1647. The “test” peptides in spots J29 and J30 are unrelated to α_1C_, ARNDQEGHILKMFPSTYVARNDQEG and WRNDQEGHILKMFPSTYVARNDQEG.

**Figure S3, related to Fig. 4.**
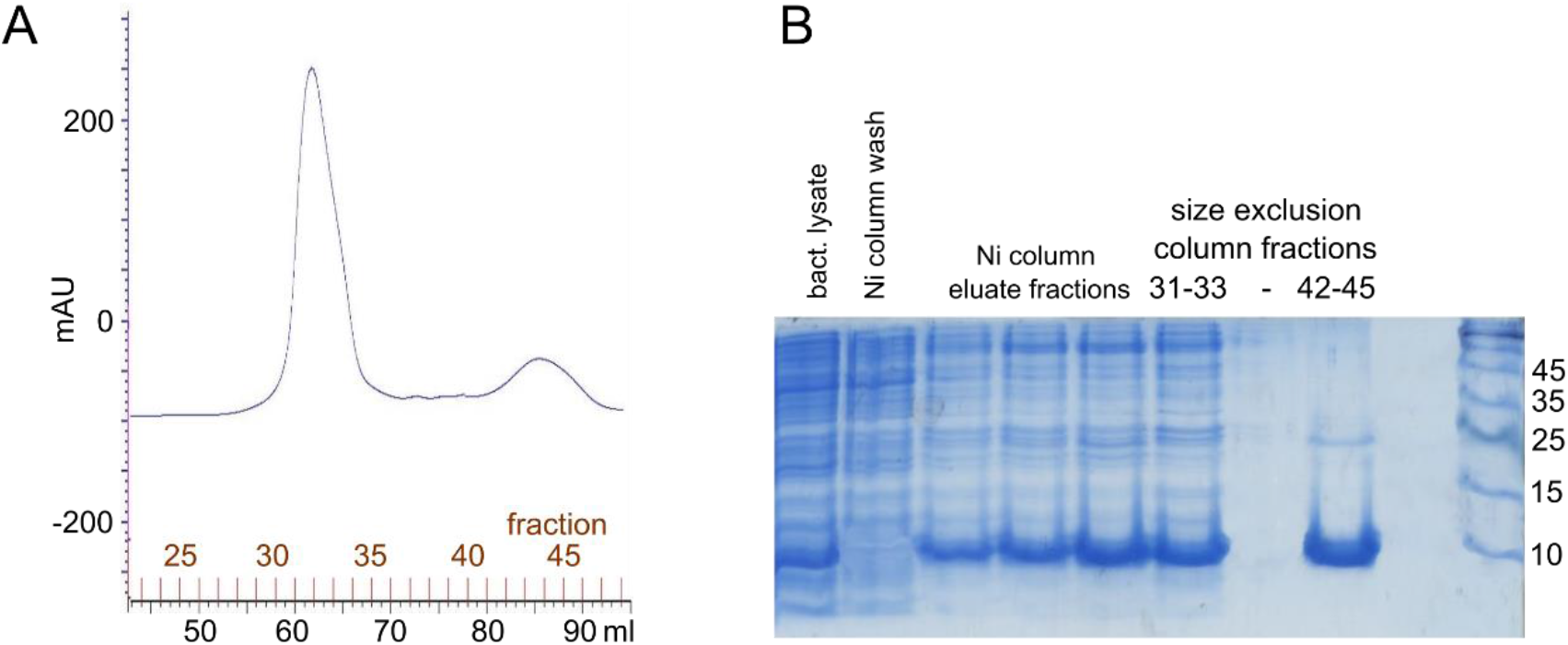
His-DCRD-Myc: purification from E. coli. A, elution profile from the size exclusion chromatography column (16/60 Superdex 75). B, analysis of the proteins obtained at the different stages of His-SUMO-PCRD preparation and purification. Coomassie stain of SDS-PAGE (12% gel). Representative of at least 3 similar purifications.

**Figure S4, related to Fig. 4.**
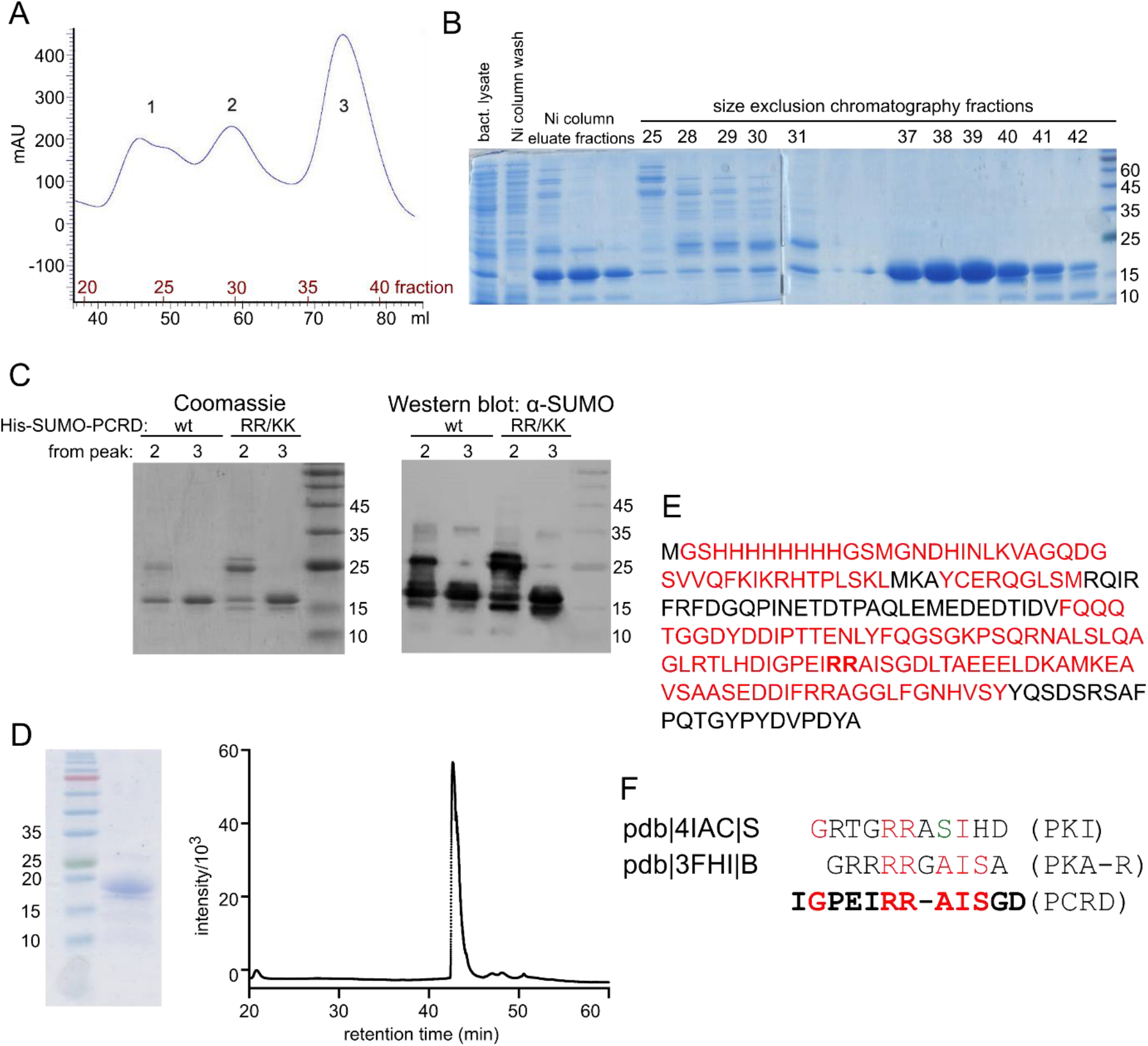
His-SUMO-PCRD: purification and analysis. *A, B,* purification of His-SUMO-PCRD from *E. coli*. A, elution profile from the size exclusion chromatography column (16/60 Superdex 75). Representative of at least 3 similar purifications. B, analysis of the proteins obtained at the different stages of His-SUMO-PCRD preparation and purification. Coomassie stain of SDS-PAGE (12% gel). *C*, Western blot with anti-SUMO antibody of purified WT and RR/KK His-Sumo-PCRD proteins (two separate preparations, the elution profiles from the Superdex column were similar to that in A). The SUMO antibody recognizes both the long and the truncated forms of His-SUMO-PCRD, indicating that the N-terminal part of the protein is not proteolysed. *D-E,* fraction 28 of His-SUMO-PCRD_trunc_ from a separate purification was analyzed by 12% SDS-PAGE and reverse phase HPLC (D), and trypsin/pepsin digestion followed by nano-LC MS/MS (E). The a.a. labeled in red color were present in the digestion products detected by the MS. *F*, alignment of PKAC-binding pseudosubstrate peptides from mouse/human PKI and mouse/bovine PKAR with the analogous segment of PCRD. Color coding (relative to PCRD): red, identical; green, similar.

**Figure S5, related to Fig. 5.**
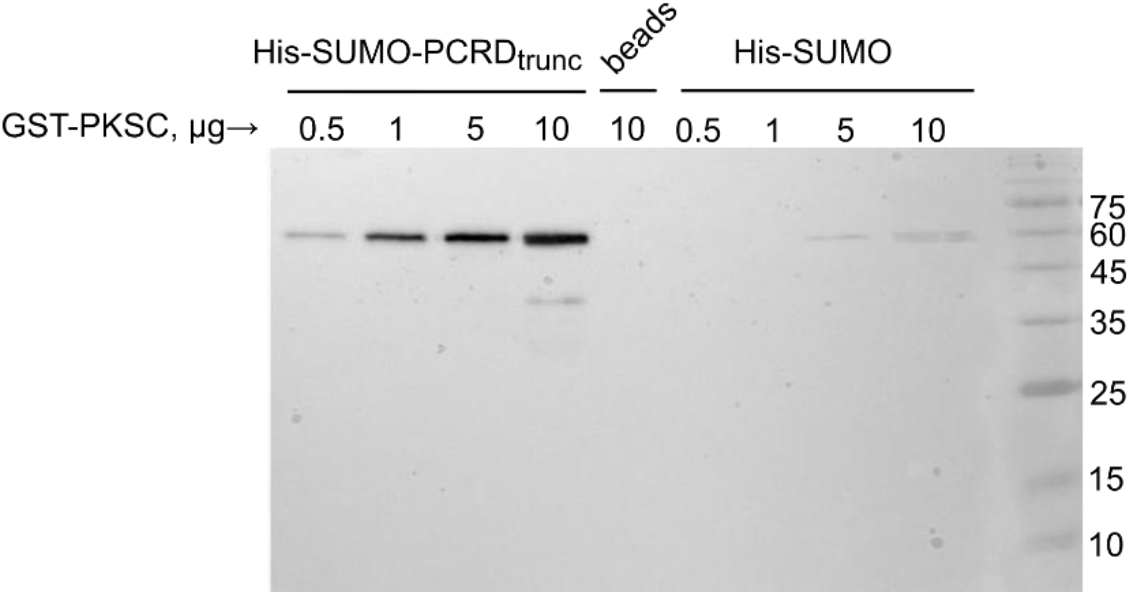
SUMO does not bind PKAC. 1 µg of purified His-SUMO-PCRD_trunc_ or His-SUMO were used to pull down the indicated amounts of GST-PKAC on Ni-NTA beads. The reaction volume was 300 µl. The image shows Western blot of the eluted proteins, detected with the PKA antibody. Representative of 2 experiments.

**Figure S6.**
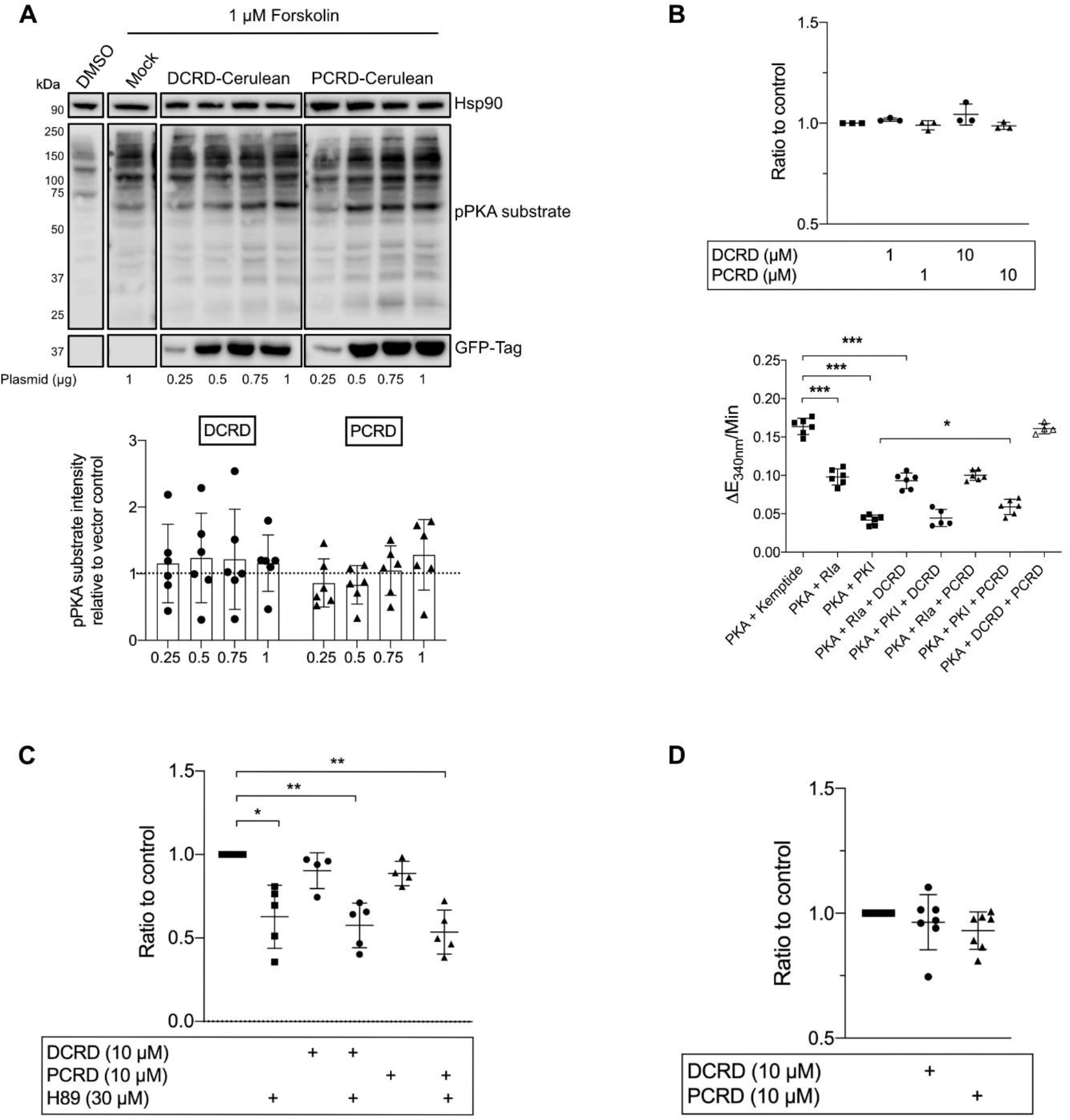
PCRD and DCRD do not alter the catalytic activity of PKAC. *A,* the indicated amounts of DCRD-and PCRD-Cerulean plasmids were expressed in HEK293 cells and the amount of phosphorylated PKA substrate proteins detected using anti-pPKA substrate antibody. The empty vector control (Mock) shows that the forskolin stimulation worked and induced phosphorylation of PKA substrates. The graph shows the mean of the pPKA substrate signal ± SD to the Mock control sample (n=6). *B,* in the Cook assay, 40 nM of human PKAC was incubated with the indicated concentrations of His-DCRD-Myc or His-SUMO-PCRD with 260 µM Kemptide. The upper graph shows the ratio to the PKAC and kemptide only control measurements ± SD (n=3). For the lower graph, 30 nM PKA-RIα or 30 nM PKI were added in the presence and absence of the channel fragments (10 µM) (n = 6 independent measurements, n = 4 for DCRD/PCRD combination). *C,* ADP Glo Assay. 10 µM of His-DCRD-Myc, His-SUMO-PCRD or 30 µM H89 were incubated with 20 nM bovine PKAC and the resulting amount of ADP measured as luminescence signal. The graph shows the ratio to the control sample (PKAC + kemptide) ± SD (n=4). *D,* cumulative graph of the Cook Assay (B) and ADP Glo Assay (C). For statistics, Kruskal-Wallis test with Dunn’s correction was applied. *p ≤ 0.05, ** p ≤ 0.01, *** p ≤ 0.001.

**Figure S7, related to Fig. 8.**
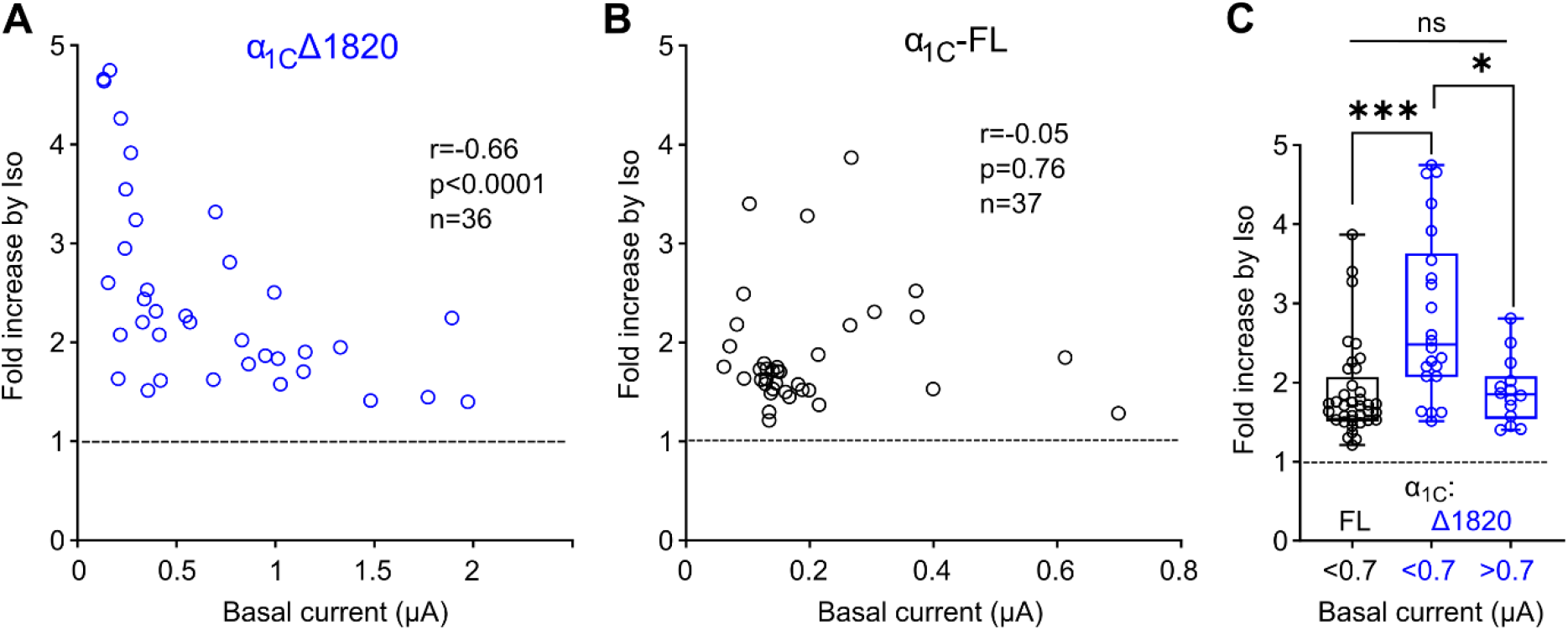
Quantitative differences in β1AR regulation of full-length and truncated Ca_V_1.2: reanalyzing the published results. The original data from Katz et al. paper (54) were analyzed in the same way as shown in Fig. 8, to seek for correlation between I_Ba_ and the extent of current increase caused by Iso (50 µM Iso were used in this work). Oocytes expressed all Ca_V_1.2 subunits (α_1C_, β2b, α2δ), rad and β1AR. *A* and *B*, correlation between basal I_Ba_ and fold increase by Iso in individual cells. Parameters of Spearman correlation analysis for the two distributions are shown in insets. Note that the amplitudes of I_Ba_ in Ca_V_1.2 with FL-α_1C_ were smaller (<0.7 µA) than for α_1C_Δ1821 (0.1-2 µA). *C*, Comparison of the differences in Iso-induced increase in I_Ba_ of α_1C_Δ1821 and α_1C_-FL for basal I_Ba_ of small-intermediate and high amplitudes. In this series of experiments large (>0.7 µA) currents were observed only with α_1C_Δ1821. Despite the smaller number of measurements, the tendency observed in experiments of Fig. 8E is also observed here: the Iso-induced increase in I_Ba_ is larger for α_1C_Δ1821 than in α_1C_-FL for smaller currents, but not for larger α_1C_Δ1821 currents.

